# Can increased prenatal exposure to thyroid hormones alter physiology and behaviour in the long-term? Insights from an experimental study in Japanese quails

**DOI:** 10.1101/2023.03.22.533811

**Authors:** Kalle Aho, Antoine Stier, Tom Sarraude, Bin-Yan Hsu, Suvi Ruuskanen

## Abstract

Maternal thyroid hormones (triiodothyronine T3 and thyroxine T4) are important regulators of embryonic development and gene expression. While maternal thyroid disorders are known to cause developmental issues in humans, variation in maternal thyroid hormones in healthy mothers have also been found to correlate with infant and child phenotypes. This suggests a potential impact of maternal thyroid hormones on offspring phenotype in an eco-evolutionary context. In chickens, prenatal thyroid hormone treatment has been shown to influence embryonic gene expression and postnatal treatment to influence imprinting and learning. However, the potential long-term effects of maternal thyroid hormones on physiology and behaviour are unclear. This study aims to investigate the long-term effects of maternal thyroid hormones on behaviour, plasma thyroid hormone levels and brain gene expression using the Japanese quail (*Coturnix japonica*) as a model. Egg hormone levels were elevated by injecting unincubated eggs with either saline (control), T3, T4 or a mixture of T3 and T4. Social motivation, boldness and fearfulness to predators were tested shortly after hatching and as adults. Plasma thyroid hormone levels and pallial expression of thyroid hormone receptor A, type 2 deiodinase, and nuclear receptor coactivator 1 were measured in adulthood. We found no evidence that elevated thyroid hormone levels in eggs affected behaviour, plasma hormone levels, or gene expression. This is the first study examining the potential long-term effects of elevated maternal thyroid hormones within the natural range. Although we found no evidence of long-term effects, other traits may still be affected and remain to be studied.

## Introduction

Maternal effects refer to the influence from maternal phenotype on their offspring’s phenotype apart from genomic inheritance (Mousseau and Fox, 1998; Moore et al., 2019; Yin et al., 2019). This influence can be mediated by hormones passed from the mother to the offspring. As the levels of maternal hormones are influenced by the maternal environment, they can transmit information about the maternal environment to the offspring. This allows the development of the offspring phenotype to be shaped by environmental factors apart from the inherited genome (Marshall and Uller, 2007; Mousseau and Fox, 1998). Potentially, if offspring environment matches with maternal environment, such hormone-mediated maternal effects can have a great adaptive potential (sensu “anticipatory maternal effects”, Marshall and Uller, 2007). Over the past decades, effects of maternally-transferred hormones on offspring phenotype have been observed in a wide range of traits across the whole animal kingdom, including physiology and behaviour (e.g. Brown et al 2014; Dantzer et al. 2013; Groothuis et al., 2019, 2005; Ruuskanen and Hsu, 2018), and can have long-lasting effects into adulthood with potential fitness consequences (e.g. metabolic rate, predator-avoidance behavior, survival, reproduction; Ruuskanen & Laaksonen 2010, 2012: Groothuis et al., 2005; Schoech et al., 2011; von Engelhardt and Groothuis, 2011). Nevertheless, a recent meta-analysis (Mentesana et al., 2025) found nearly no overall effect, in contrast with previous studies (Podmokła et al., 2018; Moore et al. 2019), suggesting that more needsto be done in order to better understand hormone-mediated maternal effects (Groothuis, Hsu, Kumar 2020).

Compared to the more extensively studied androgens and glucocorticoids (Podmokła et al., 2018), the effects of maternally transferred thyroid hormones (THs) have been less explored (Darras, 2019; Ruuskanen and Hsu, 2018). THs (triiodothyronine, T3, and thyroxine, T4) play important roles in metabolism and development, where they regulate the expression of numerous genes (Bernal, 2017; Darras, 2019; Moog et al., 2017). TH action is regulated by several components of the thyroid system. TH receptors (THRA and THRB) and coactivators control the transcription of TH-responsive genes by forming a complex with the active form of the hormone, T3 (Brent, 2012). The intracellular availability of T3 is regulated by several transmembrane TH transporters (e.g. MCT8, OATP1c1) that differentially transport either T4 or T3 (Groeneweg et al., 2020). Inside the cell, T3 availability can be further regulated by deiodinase enzymes that either convert T4 to T3 (deiodinase 2, D2) or inactivate both T3 and T4 (deiodinase 3, D3) (Gereben et al., 2008). Interestingly, the expression of these regulatory components (TH receptors, transporters and deiodinases) begins as early as embryonic day 4 in chickens (Flamant and Samarut, 1998) and embryonic day 1 in passerines (Ruuskanen et al. 2022), allowing the embryo to regulate TH levels in an organ and cell-specific manner (Darras, 2019; Darras et al., 2011; Flamant and Samarut, 1998; Ruuskanen et al., 2022; Van Herck et al., 2012) , while the development of the thyroid gland occurs later in development, around mid-incubation (Forhead and Fowden, 2014; McNichols and McNabb, 1988; Porazzi et al., 2009). This suggests that these molecular components are in place to regulate maternally-transferred hormones. Indeed, in almost all vertebrates studied so far, proper embryonic development depends on maternally transferred THs (Brown et al., 2014; Darras, 2019; Moog et al., 2017; Patel et al., 2011).

In mammals, maternal THs are transferred across the placenta, whereas in egg-laying animals such as birds, maternal THs are deposited in the egg yolk. In birds, high natural variation in egg TH levels has been observed, up to 5-fold in magnitude (Ruuskanen and Hsu, 2018). This variation may be due to environmental factors such as food availability or temperature (Hsu et al., 2016; Ruuskanen et al., 2016c), and has been reported at both the inter- and intra-individual levels (Hsu et al., 2016, 2019b; Ruuskanen et al., 2016a, 2016c). As THs are important in development (Darras et al., 2009; Decuypere et al., 2005; McNabb, 2007; Wilson and McNabb, 1997), and as increasing yolk TH levels has been found to increase embryonic TH levels (Van Herck et al., 2012), high variation in egg TH levels could therefore be a source of offspring phenotypic variation.

To date, most of what is known about phenotypic differences due to maternal THs comes mainly from biomedical studies (Moog et al., 2017). However, studies using pathological TH concentrations do not provide relevant information on the ecology or evolution of maternal THs. Therefore, the effects of maternal THs within the natural range need to be studied to understand whether variation has effects on traits affecting offspring fitness. In addition, experiments manipulating maternal THs in mammals would inevitably interfere with maternal physiology, making it difficult to attribute results solely to maternal THs. Egg-laying animals, such as birds, have therefore become common models for studying the functions of maternal hormones since maternal THs deposited in eggs can be manipulated directly and independently. So far, elevated TH levels in the egg within the natural range have been found to affect hatching success, offspring growth, chick levels of T3 and telomere length in altricial bird species (Hsu et al., 2023a, 2019a, 2017; Ruuskanen et al., 2016b; Sarraude et al., 2020; Stier et al., 2020). Nevertheless, some of the reported effects appear to be transient during the early post-hatching period (Hsu et al., 2019a). Whether the effects of maternal THs can be long-lasting and potentially influence offspring fitness remains unclear. In terms of precocial bird species, previous studies studying maternal THs were scarce and typically applied supra-physiological doses (e.g., Wilson and McNabb, 1997; van Herck et al., 2012).

To investigate the potential long-term effects of variation in maternal THs, we experimentally elevated egg TH levels within the natural range in Japanese quail (*Coturnix japonica*). With these experimental quails, we have previously reported a higher hatching success caused by the elevated yolk TH levels, but no major effects on morphology or physiology (Sarraude et al., 2020). Now we further studied changes in chick and adult behaviour and adult plasma TH levels. In addition, we studied brain gene expression at a later age. While increasing maternal TH levels beyond normal physiological range (maternal hyperthyroidism) has been shown to affect behaviour, for example both decreasing and increasing anxiety (Laureano-Melo et al., 2019; Zhang et al., 2010, 2008), it is not known whether increasing maternal THs within the natural range affects similar or linked behaviors. Also, as postnatal THs have been shown to influence filial imprinting and learning behaviour (Yamaguchi et al., 2012), it is interesting to study whether maternal THs, acting earlier in development, may also have similar behavioural effects either in chicks or in adulthood. The underlying mechanistic links between thyroid hormones and anxiety are expected to link TH-induced changes in early brain development (in humans, for example altered hippocampal neurotransmitter release or other neurochemical changes, even changes in neurogenesis, summarized in Laureano-Melo et al. 2019 Horm Behav). We therefore studied traits related to anxiety (Kumar et al. 2013) in our animal model, using existing validated protocols testing boldness/fearfulness and social stress (Kumar et al. 2013, Forkman et al. 2007). Such traits related to boldness and fearfulness are very relevant in any animal model as they have clear links to for example foraging success and risk of predation. Potential behavioural effects were investigated in both chicks and adults, allowing for a broader timescale. Due to the lack of studies with natural levels of elevated maternal THs and inconsistent results from studies on maternal hyperthyroidism and offspring anxiety (Laureano-Melo et al., 2019; Zhang et al., 2010, 2008), the direction of potential behavioural changes is difficult to predict. In addition, plasma TH levels were measured in adults, and changes in these would indicate long lasting effects on TH function.

While plasma TH levels have been reported to be affected by elevated egg TH levels (within the natural range) in a sex-dependent manner in nestling rock pigeons (Hsu et al., 2017), some more recent studies in wild passerines found no clear effects of higher natural levels of egg THs on nestling plasma THs (Hsu et al., 2023b, 2020). In terms of gene expression, previous studies have examined the effect of the natural variation in maternal THs on brain gene expression only in bird embryos or young chicks (Van Herck et al., 2015, 2012; Yamaguchi et al., 2012) focusing on developmental effects. However, the potential long-term effects on TH action in the adult brain remain unknown. Here we studied several genes involved in TH function: THRA (thyroid hormone receptor A), NCOA1 (nuclear receptor coactivator 1) and DIO2 (type 2 deiodinase) in the pallium of adults. Expression of these genes has been shown to be influenced by THs in other species (Misiti et al., 1998; Walpita et al., 2007). As these genes are part of the cellular pathway of TH action, changes in their expression could indicate long-term changes in the regulation of TH action in the brain, and consequently, brain function.

## Materials and methods

### Materials

This study was conducted in accordance with the Finnish legislation and approved by the Finnish Animal Ethics Committee (Eläinkoelautakunta; ESAVI /1018 /04.10.07 /2016). Japanese quail (*Coturnix japonica*), a commonly used laboratory model with a reference genome, was used. This study was done with same individuals previously used in Sarraude et al. (2020).

### Parental generation and egg collection

The parental generation was composed of adult quails that were supplied by private breeders in Finland and kept in two acclimatised rooms. Twenty-four breeding pairs were formed by pairing birds from different breeders. Individuals were identified using metal leg rings. The floor was covered with 3–5 cm of sawdust bedding, and a hiding place, sand and calcium grit were provided. Each pair was housed in an indoor aviary divided into pens with a floor area of 1 m^2^. The ambient temperature was maintained at 20°C, and the photoperiod was set to 16 light hours (06:00 am-22:00pm, +3UCT) and 8 dark hours. The birds were fed ad libitum (Poultry Complete Feed, “Kanan Paras Täysrehu”, Hankkija, Finland), and their water was replaced daily. Pairs were monitored every morning to collect eggs for 7 days (longer storage time is known to decrease hatchability). The eggs were marked with a non-toxic marker, weighed, and stored in a climate-controlled chamber at 15°C and 50% relative humidity. On the final day of the collection period, a solution was injected into a total of 4–8 eggs per pair (the variation was due to not all females producing an egg every day, average = 7, standard deviation = 1)

### Treatment of eggs, rearing of chicks and collection of samples

We manipulated egg TH levels by injecting hormones directly into the eggs prior to embryonic development at a dose that mimicked increased maternal transfer (see details in Sarraude et al., 2020). Eggs from each breeding pair were randomly assigned to four treatment groups: T3, T4, T3 + T4 and control (0.9% saline), 24 clutches for all groups except T3 that had 22 clutches. We used saline control following previous studies on injecting THs into eggs (van Herck et al. 2015), and because THs are not very well soluble in oil. Eggs laid by the same pair were always evenly distributed among all treatment groups. The injected hormone solutions were prepared from crystallised powder (T3: CAS number 6893-02-3, Sigma-Aldrich, St. Louis, MO, USA and T4: CAS number 51-48-9, Sigma-Aldrich, St. Louis, MO, USA). The amount of hormone injected (T4: 8.9 ng/egg, equivalent to 1.79 pg/mg yolk; T3: 4.7 ng/egg, equivalent to 1.24 pg/mg yolk) was based on previous measurements in the parental generation (mean ± SD: T4, 15.3 ± 4.43 ng /egg; T3, 7.58 ± 2.32 ng /egg) and the aim was to increase hormone contents in the yolk by two standard deviations (as recommended by Podmokla et al., 2018). We were unsure how much of T4 is converted to T3 and therefore used 2SD of both hormones for the T3T4 group. The injection protocol has been described in detail in Sarraude et al. (2020). Briefly, each egg was injected with 50 µl of either the respective hormone solution (T3, T4 or T3+T4, in 0.9% saline) or 0.9% saline (control, CO), delivered by using 27G insulin syringes (BD Microlance^TM^). After sealing the hole with sterile plaster (OPSITE Flexigrid, Smith & Nephew), the eggs were incubated artificially (37.8 °C and 55% relative humidity). A total of 158 eggs were treated (39 T3, 39 T4, 40 T3+T4 and 40 CO), of which 66 chicks hatched (15 T3, 20 T4, 21 T3 + T4 and 10 CO), with sex distributions of chicks surviving to adulthood (females /males): 7/4 for T3, 10/8 for T4, 9/12 for T3 + T4 and 3/4 for CO. Breeding pairs/pairs of siblings per treatment: CO 7/0, T3 9/2, T3T4 15/4 and T4 13/2. Sex was determined at maturity by examining the presence of the cloacal gland. The overall hatching rate (∼40%) was consistent with previous studies in domestic quail (Okuliarová et al., 2007).

Twelve hours post-hatching, the chicks were marked with a unique combination of coloured rings and nail coding, and transferred to two cages (each with a floor area of 1 m2 and a height of 30 cm). The cages contained approximately 30 chicks, sex and treatments mixed. The chicks were provided with heating mats and lamps as supplementary heat sources for the first two weeks. The chicks were fed with sieved commercial poultry feed (“Punaheltta paras poikanen”, Hankkija, Finland) and provided with calcium and bathing sand. Two weeks after hatching, the chicks were separated into four 1 m2 cages (ca. 30 cm high) of approximately 16 individuals. Approximately three weeks after hatching, the coloured rings were replaced with unique metal rings. At week 4 after hatching, the birds were transferred to eight pens of 1 m2 floor area (average of 7.1 birds/pen, range = 4–9), under the same conditions as the parents. At around the age of sexual maturity (ca. 6–8 weeks after hatching), the birds were separated by sex in twelve 1 m2 pens (average of 4.8 birds/pen, range = 4–5). Photoperiod was 16L:8D for the first 7 months and 8L:16D thereafter. At approximately 4.5 months of age, we collected blood samples (200 µl) from the brachial vein, separated plasma, and stored it at -80° C. As Japanese quails have reached adulthood at 4.5 months, this time point would have already shown if there are long-lasting, organizational effects of prenatal THs effects into adulthood. All birds were euthanised by decapitation at 10 months of age. After decapitation, brain tissue was collected for the gene expression analysis from the pallium (average 44.0 mg, std 2.8 mg, taken at the surface of the region) in PBS buffer, snap frozen in liquid nitrogen and stored at -80° C. While differences in photoperiod limit comparisons between gene expression results and other results, our aim was to study whether there are persistent effects of prenatal THs independent of any environmental factors that individuals may face later in life, for which our experimental design is sufficient.

### Behavioural experiments

To assess the potential effects of maternal THs on behaviour, we conducted a series of behavioural tests (open arena, emergence and tonic immobility tests) at different life stages. We followed previously published protocols that have been used extensively with many different model organisms. The tests were conducted in separate rooms where the birds had no visual or auditory contact with other individuals. The exception was the open arena test, where the subject was able to hear the vocalisations of another bird being tested at the same time in another room. All experiments were recorded, and the videos were analysed blindly of the treatment. All videos of the same test at both ages (chick/adult) were analysed by one person, except for the open arena test, where one person analysed chick videos and another person analysed adult videos. This may have confounded the age difference in behavior scores with the observer effect. Since the videos from different treatments at each age were analysed by the same person, the fact that chick and adult videos were analyzed by different people should not confound the effect of prenatal TH exposure.

The open arena test measured social motivation/stress and boldnes/fearfulness and it followed a setup developed by Forkman et al. (2007). The test was performed once on both sexes at three days (F/M/undetermined: CO 3/3/1, T3 6/3/2, T3T4 8/12, T4 8/8) and once approximately 4.5 months of age (F/M: CO 3/4, T3 8/3, T3T4 9/12, T4 9/8). In the test, the bird was placed in a circular illuminated arena and the movement duration and the number of escape attempts (attempt to jump/fly over the wall) were measured from videos for 10 minutes (from the time the researcher left the room). Only 10 of the 56 adults (1 CO, 2 T3, 2 T4 and 5 T3T4) attempted to escape, so adult escape attempts were not analysed. Subjects who moved less and made fewer escape attempts were interpreted as less bold and less socially motivated.

The emergence test measured social motivation and boldness/fearfulness (especially the timidity aspect, see e.g Davis et al. 2008). The test was performed on both sexes, twice on 6-9d old chicks (as there is less data on repeatability, Forkman et al. 2007; F/M/undetermined: CO 3/4/1, T3 7/4/1 T3T4 9/12, T4 10/8) and once on 4.5 months old adults (F/M: CO 3/4, T3 7/4, T3T4 8/11, T4 9/7) . The test followed a setup developed by Forkman et al. (2007), in which a bird was placed in a dark chamber connected to another, bright chamber. One minute after the subject was placed in the dark chamber, a door connecting the chambers was removed. After removing the door, the researcher left the observation room and the time it took for the subject to completely exit the dark chamber was measured. Subjects who left the dark chamber more quickly were interpreted as bolder and more socially motivated. The test lasted five minutes. Whether the subject remained in the dark chamber (0) or left it (1) during the test was also recorded.

The tonic immobility test (Mills and Faure, 1991) measured fearfulness to predators (Forkman et al 2007). The test was performed on once both sexes at 12 days (F/M/undetermined: CO 3/4, T3 7/4/1, T3T4 9/12, T4 10/8) and once approximately 4.5 months of age (F/M: CO 3/4, T3 7/4, T3T4 8/12, T4 9/7). To induce immobility, the bird was placed on its back in the centre of a V-shaped test apparatus and restrained by hand for 15 seconds, after which the hand was removed, and the time taken for the bird to resume a standing posture was measured. If the subject resumed the standing posture in less than 10 seconds, this was recorded as a failed induction and the induction procedure was repeated. Four adult subjects who failed five times were excluded. If the subject remained stationary for more than 10 seconds, then the time to recovery was measured. The longer a subject took to recover, the more fearful it was interpreted. The test lasted 10 minutes. Five subjects (1 chick and 4 adults) did not recover for 10 minutes and were excluded from the analysis (approximately 4% of all tests).

### Plasma thyroid hormone levels

To test whether elevated maternal THs affect plasma TH levels later in life (4.5 months of age), we extracted THs from plasma samples and measured them using a previously validated LC-MS /MS protocol (Ruuskanen et al., 2018, where CV <10%). Due to practical limitations (*i.e.* volume of plasma available and haemolysis in some samples), the sample size was reduced to 48 individuals (F/M: T3 6/4, T4 8/8, T3T4 6/9 and 3/4 in the control). Hormone concentrations are expressed in pmol/ml plasma. The repeatability of the TH measurements, was validated in Ruuskanen et al. (2018) with CV% less than 10% (3.5-9.7%, depending on the hormone and sample type)

### Gene expression

To test for potential long-term changes in the expression of TH related genes (*DIO2, DIO3, THRA*, and *NCOA1*), we measured gene expression from pallium samples (10 months old). RNA was extracted from the homogenised pallium samples using the NucleoSpin RNA Plus kit (MACHEREY-NAGEL). Since the sample size of the control group was the main factor limiting statistical power, we decided to analyse all control individuals and only an equivalent number of individuals (randomly selected) in the three treatment groups (N = 31, 3F/4M in control and 4F/4M per other treatment groups). Extraction was done from 50 µl of the homogenised brain sample according to the manufacturer’s instructions, except that the gDNA removal column was centrifuged longer (1 min, 11000 g) and the RNA was eluted once with 40 µl of Rnase-free water. RNA concentration and purity were quantified by optical density, and all samples not meeting our quality criteria (i.e. RNA concentration > 30 ng/μl and 260/280 > 1.80) were excluded from further analysis. RNA integrity was checked using an E-Gel 2% electrophoresis system (Invitrogen), and the ribosomal RNA 18S vs. 28S band intensity and was deemed satisfactory (the visual inspection revealed the expected two bands of rRNA with 28S band being > 2 times more intense than 18S band). Samples were stored at -80° C for 2 weeks prior to cDNA synthesis. cDNA synthesis was performed using the SensiFAST ™ cDNA Synthesis Kit (Bioline) according to the manufacturer’s instructions. The reaction was performed using 500 ng of RNA as the starting material. A control (noRT) was prepared without reverse transcriptase enzyme. The cDNA was diluted with water to a concentration of 1.2 ng/µl and stored at -20° C. The 1.2ng/uL was calculated from RNA pre-RT, and slight variations in cDNA starting quantity (due to differences in RT efficiency) are controlled for using the reference genes when calculating relative expression.

*GAPDH* and *PGK* were used as control genes, using primers previously designed and validated in Japanese quail (Vitorino Carvalho et al., 2019). Stability of expression between experimental groups was further verified using geNorm software (geNorm M < 0.5, geNorm V < 0.15), and thus the geometric mean of these two genes was used as our normalization factor. Primers for the target genes were designed using the Primer-BLAST tool (NCBI). The primer pairs (Table 1) were designed to have at least one intron between them. The suitability of the primers in qPCR was tested prior to the actual analysis using different primer concentrations and different amounts of cDNA. The tests confirmed the sufficient amplification efficiency of the primers (between 90 and 110% based on the standard curve) and the formation of only one product of the expected size based on both the presence of a single narrow peak in the melting curve, and the presence of a single band of the expected size on the agarose gel. *DIO3* had extremely poor amplification, meaning that we could not accurately quantify it, and was excluded from the analysis.

qPCR assays were performed on a Mic qPCR (Bio Molecular Systems) in a final volume of 12 µL containing 6 ng of cDNA, 800 nM of forward and reverse primers and SensiFAST SYBR Lo-ROX (Bioline). We ran gene-specific qPCR plates and each sample was analysed in duplicate. Two plates were run for each gene and experimental groups and sex were always balanced within each plate. We used a cDNA reference sample (*i.e.* ratio = 1), which was a pool of 10 different individuals on each plate. We used a two-step cycling with the following conditions: 2 min at 95°C; then 45 cycles of 5s at 95°C followed by 20s at 60°C (fluorescence reading) for all reactions, as recommended by manufacturer. The expression of each gene was calculated as (1+Ef_Target_)^ΔCq(Target)^/geometric mean [(1+Ef_GAPDH_)^ΔCq^^GAPDH^; (1+Ef_PGK1_)^ΔCq^^PGK1^], where Ef is the amplification’s efficiency and ΔCq is the difference between the Cq-values of the reference sample and the sample of interest. Amplification in the no-RT controls never occurred until at least 8 cycles after the lower Cq sample, and thus contamination by genomic DNA could not interfere with our results. Technical precision and repeatability were generally good (Table 1), but a few samples with a CV% of duplicates (based on the final ratio) greater than 15% were excluded from the analysis (final sample size: N_DIO2_ = 27, N_NCOA1_ = 28, N_THRa_ = 26)

### Statistical analysis

All statistical analyses were performed with R software (version 3.6.2), using the packages lme4 version 1.1-23 (Bates et al., 2015) and emmeans version 1.6.0 (Lenth, 2021). Normality and homoscedasticity of the residual distributions were checked visually (quantile-quantile and residuals against fitted values plots)with additionally confirmation by Shapiro-Wilk test of normality. For the behavioural analysis, recovery time (tonic immobility) and exit time (emergence test), were natural log-transformed to ensure that the residuals were normally distributed. Possible outliers were tested with R using the *simulateResiduals* function in DHARMa package (Hartig 2024).

Linear mixed models (LMM, *lmer* function in lme4) were used to analyse time to exit in the emergence test, movement duration and number of escape attempts in the open arena test, recovery time in the tonic immobility test, plasma TH concentrations and qPCR results. Generalised linear mixed model (GLMM, g*lmer* function in lme4) with a binomial error distribution was used to analyse the proportion of individuals leaving the dark chamber in the emergence test. Fixed effects were treatment, sex, and test number/age (in repeated tests). Interactions between sex and treatment, and between treatment and age (in repeated tests), were also analysed. Random effects were individual ID (in repeated tests) and parental ID (to account for relatedness between siblings in all experiments). F- and p-values for LMMs were obtained using the Kenward-Roger approximation on the denominator degree of freedom (ddf), implemented in the package *lmerTest* (*anova* function) (Kuznetsova et al., 2017). For binomial regression models, main effects and interactions were tested by comparing the model with and without the factor. For all models, we removed non-significant (p > 0.05) interactions to ensure correct interpretation on the main effects. Post-hoc tests were performed using Tukey tests (*emmeans* function from emmeans package, Lenth, 2021) where appropriate.

## Results

### Effects of prenatal thyroid hormone supplementation on behaviour

In general, no statistically significant differences in chick or adult behavioural traits were observed between the egg TH manipulation groups (T3, T4, T3 + T4, control). Elevated THs had no significant effect on social motivation or boldness (Table 2, Fig. 1-2), nor on fearfulness to predators (Table 2, Fig. 3). A statistically significant interaction between treatment and age was found in the emergence test (F = 3.58, ddf = 67.2, p = 0.018, Fig. 2): In all treatments except the control group (Tukey t=0.59, p=0.56), chick exit times were significantly longer than those of adults (all t > 3.21, p < 0.002) while no differences between treatments were found within each age-category (all t < 2.48, all p> 0.07). There was also a statistically significant interaction between treatment and sex in the tonic immobility test (F = 2.92, ddf = 37.5, p = 0.047, Fig. 3): In males, the recovery time was longer in the T4-group (marginal mean ± SE: 177.1 ± 47.8 s) than in the T3-group (marginal mean ± SE: 56.1 ± 20.8 s, Tukey t = 2.71, p = 0.047), whereas no differences between treatments were observed in females (all t < 2.29, all p >0.12). In the T4-group, the recovery time of males (marginal mean ± SE: 177.1 ± 47.8 s) was longer than that of females (marginal mean ± SE: 85.2 ± 19.9 s) (t=2.16, p= 0.037), while no sex differences were observed within the other treatments (all t<1.80, all p>0.079).

**Figure 1.**
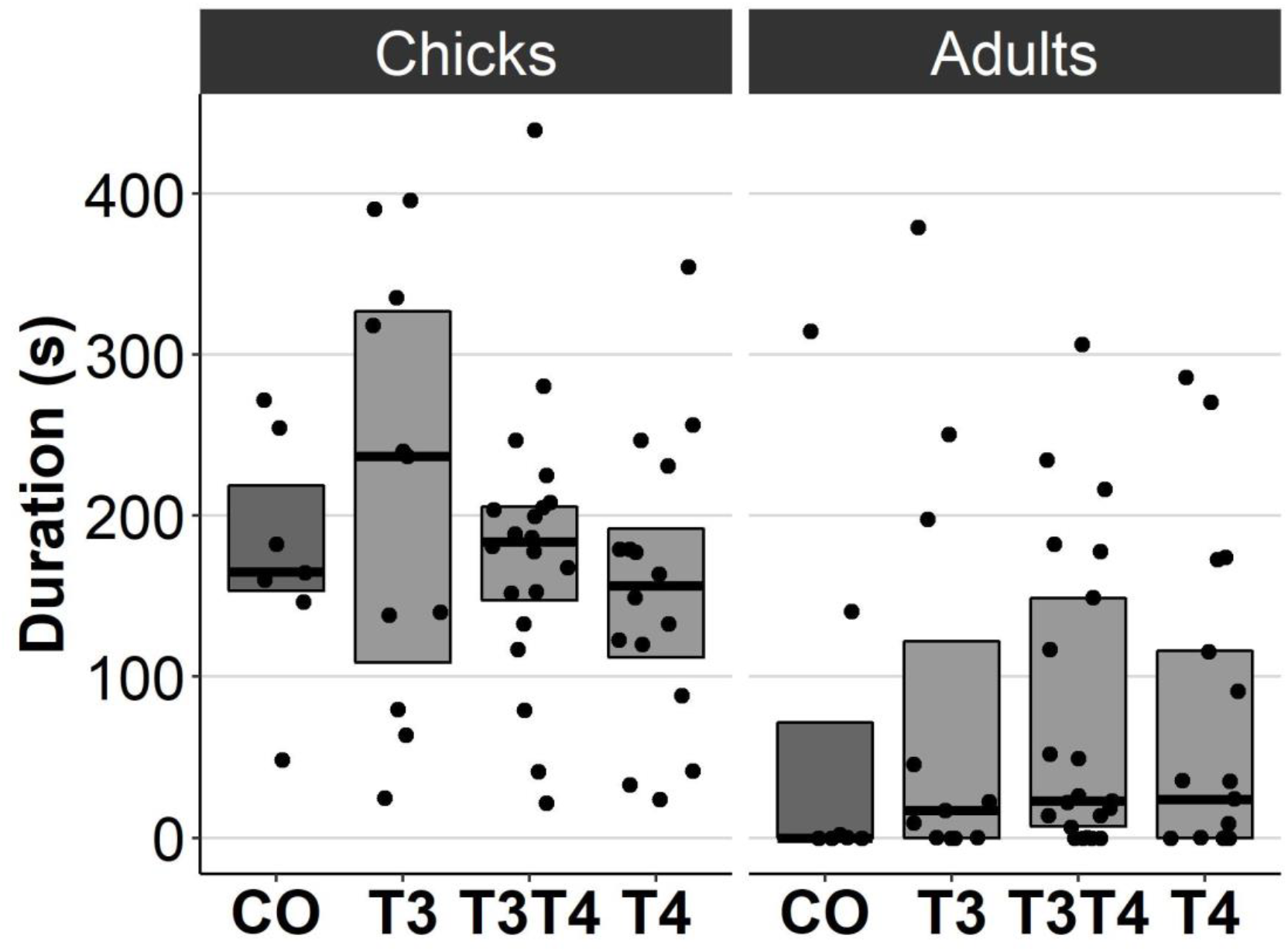
Movement duration for chicks (3 d) and adults (4.5 months) in the open arena test for different prenatal thyroid hormone treatment groups. CO = control, 50 µl 0.9% saline; T3: 4.7 ng/egg in 50 µl 0.9% saline, equivalent to 1.24 pg/mg yolk; T4: 8.9 ng/egg in 50 µl 0.9% saline, equivalent to 1.79 pg/mg yolk. The lines represent the medians of the treatments, and the bars represent the interquartile range of the original observations. Sample size (chicks/adults): CO 7/7, T3 11/11, T3T4 20/21 and T4 16/17.

**Figure 2.**
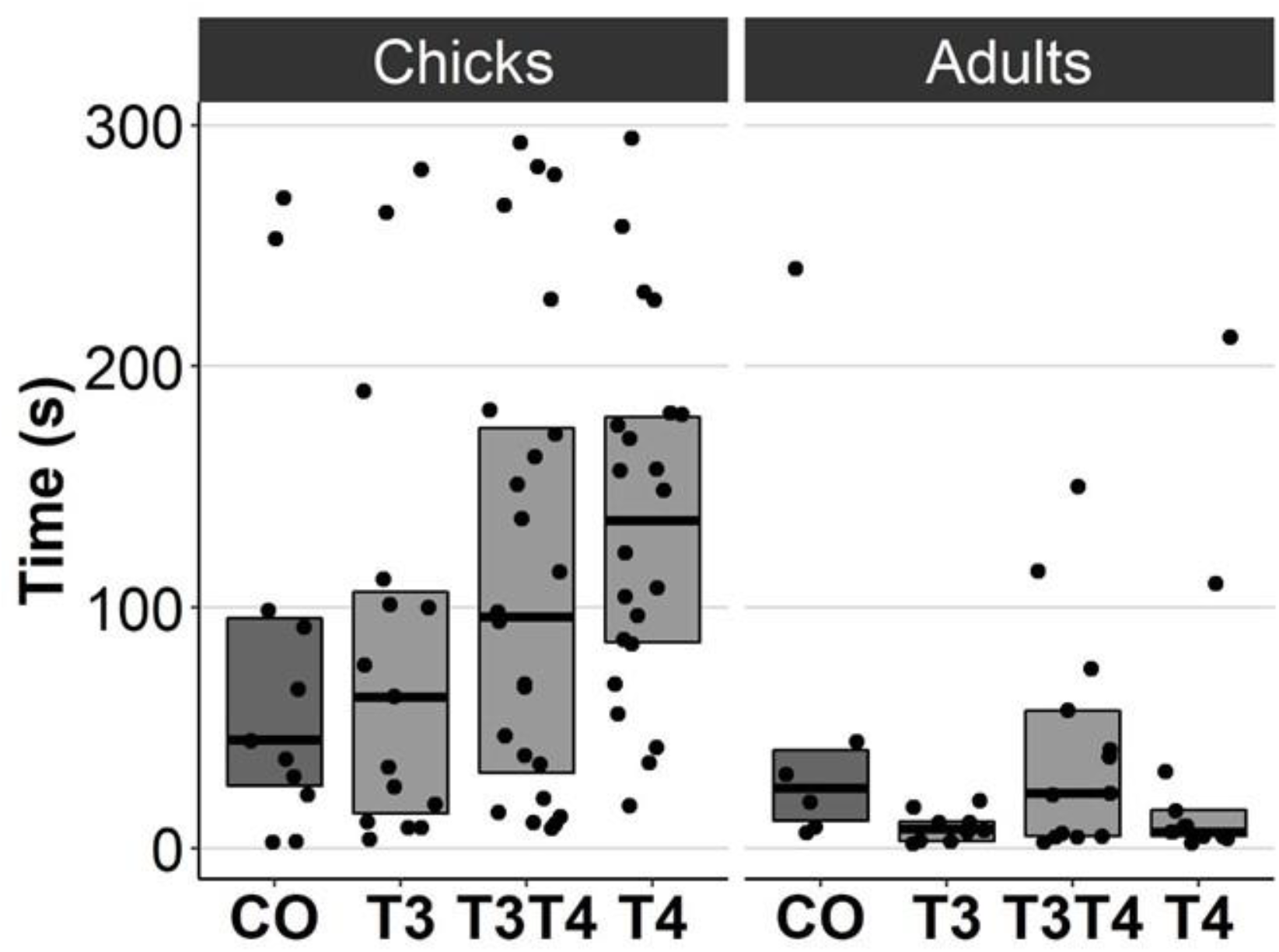
Time to exit the dark chamber for chicks (6-9 d) and adults (4.5 months) in the emergence test for different prenatal thyroid hormone treatment groups. CO = control, 50 µl 0.9% saline; T3: 4.7 ng/egg in 50 µl 0.9% saline, equivalent to 1.24 pg/mg yolk; T4: 8.9 ng/egg in 50 µl 0.9% saline, equivalent to 1.79 pg/mg yolk. The test was performed twice for chicks and once for adults. Results for both chick studies are shown. The lines represent the medians of the treatments, and the bars represent the interquartile range of the original observations. Sample size (chicks/adults): CO 11/6, T3 15/9, T3T4 24/13 and T4 22/13.

**Figure 3.**
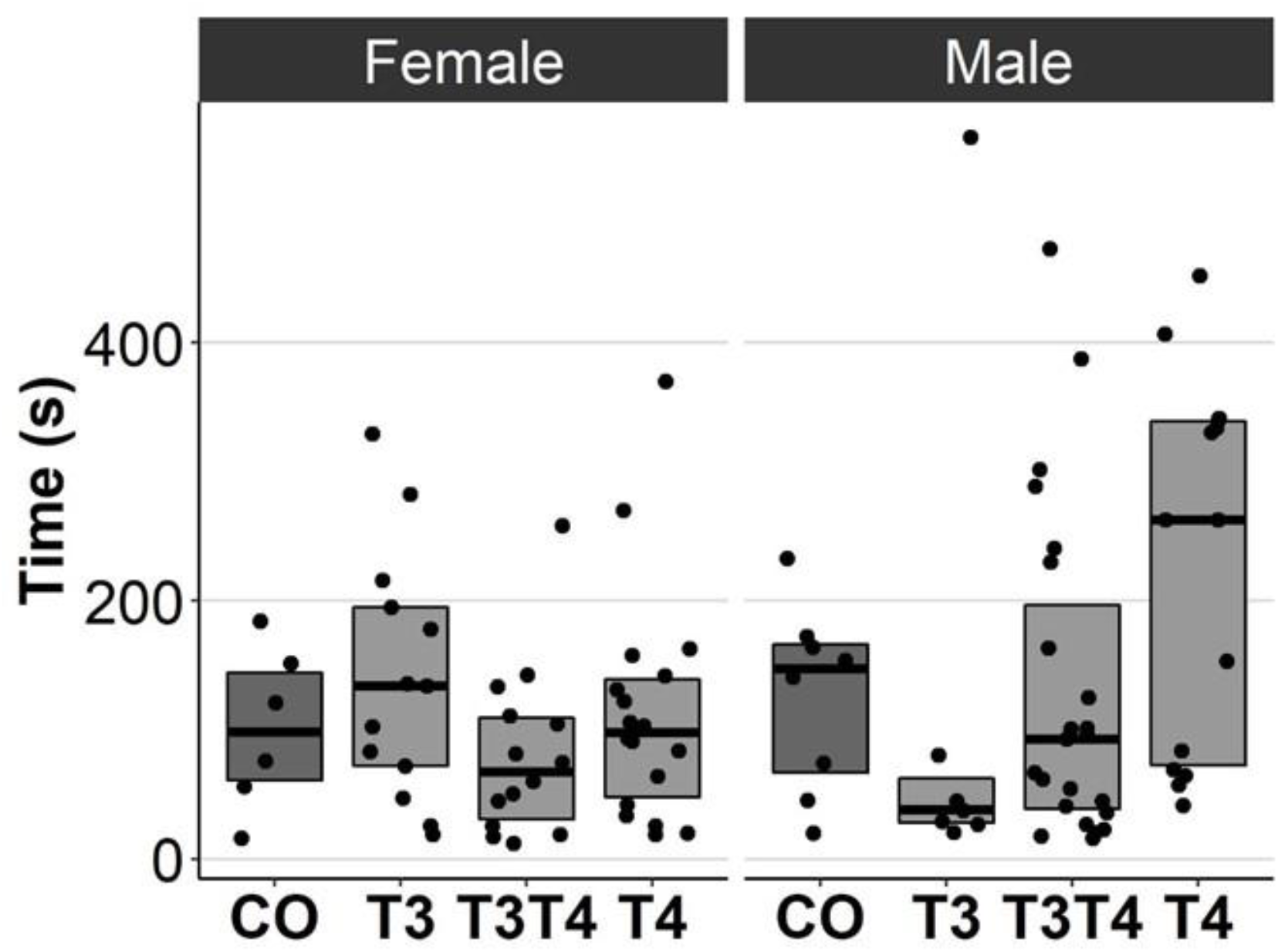
Time to resume standing position for females and males, chicks (12 d) and adults (4.5 months) combined, in the tonic immobility test for different prenatal thyroid hormone treatment groups. CO = control, 50 µl 0.9% saline; T3: 4.7 ng/egg in 50 µl 0.9% saline, equivalent to 1.24 pg/mg yolk; T4: 8.9 ng/egg in 50 µl 0.9% saline, equivalent to 1.79 pg/mg yolk. The lines represent the medians of the treatments, and the bars represent the interquartile range of the original observations. Sample size (female/male: CO 6/8, T3 13/7, T3T4 14/23 and T4 18/14.

### Effects of prenatal thyroid hormone supplementation on plasma thyroid hormone levels

Increasing TH levels in eggs did not significantly affect plasma TH levels in 4.5-month-old birds (T3: F = 0.13, ddf = 37.7, p = 0.94, T4: F = 2.1, ddf = 41.5, p = 0.12, Fig. 4). Treatment also had no sex-dependent effect on plasma T3 or T4 (treatment-sex interactions, p > 0.36).

**Figure 4.**
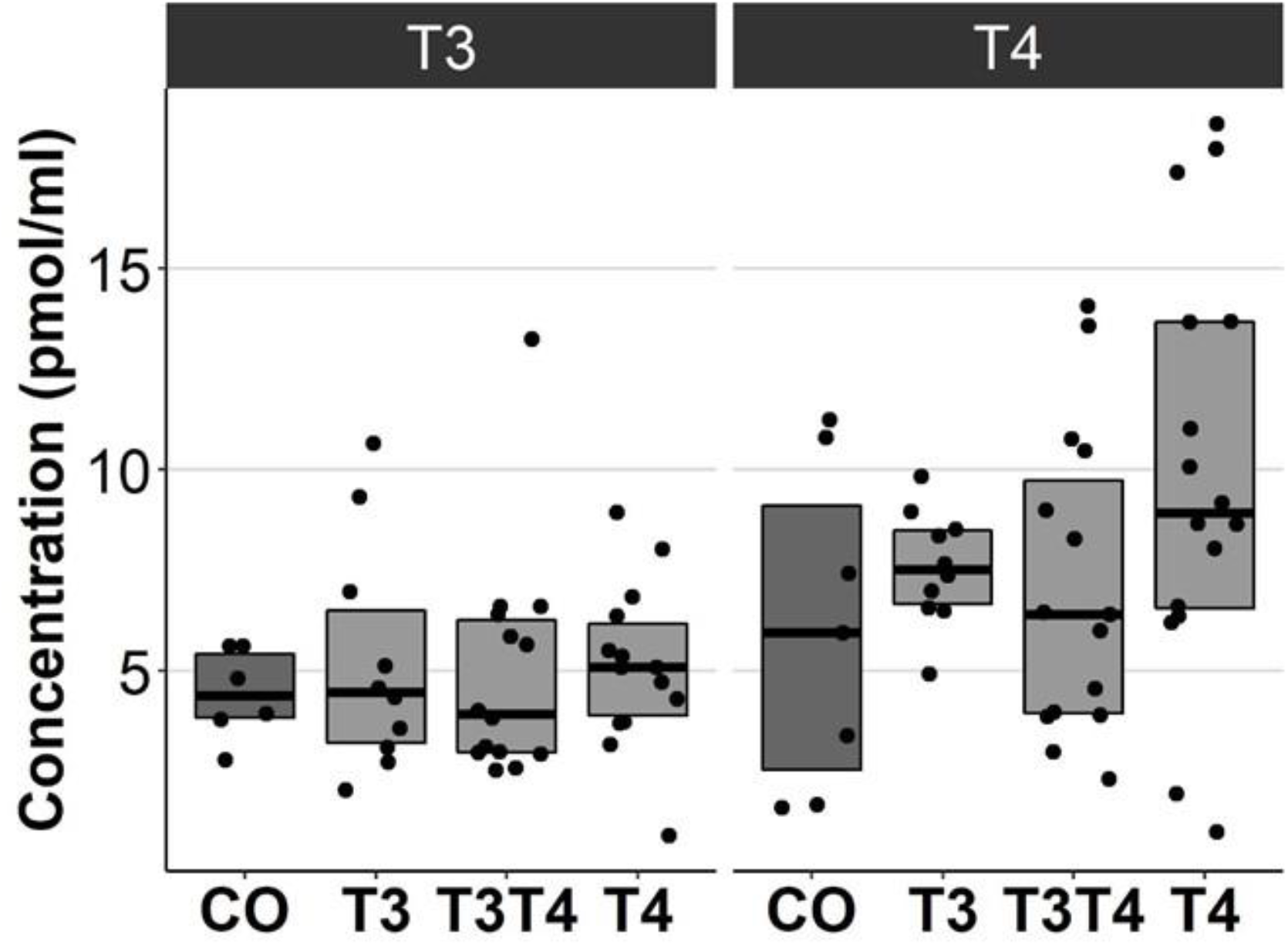
Plasma T3 and T4 concentrations in 4.5-month-old birds between different prenatal thyroid hormone treatment groups. CO = control, 50 µl 0.9% saline; T3: 4.7 ng/egg in 50 µl 0.9% saline, equivalent to 1.24 pg/mg yolk; T4: 8.9 ng/egg in 50 µl 0.9% saline, equivalent to 1.79 pg/mg yolk. The lines represent the medians of the treatments, and the bars represent the interquartile range of the original observations. Sample size (T3/T4 concentrations): CO 6/7, T3 10/10, T3T4 14/15 and T4 14/16.

### Effects of prenatal thyroid hormone supplementation on pallial gene expression

Increasing TH levels in eggs did not affect pallial gene expression of TH-related genes in 10-month-old birds (*DIO2*: F = 2.49, ddf = 17.3, p = 0.09, *NCOA1*: F = 0.092, ddf = 21.1, p = 0.96, *THRA*: F = 0.47, ddf = 18.9, p = 0.71, Fig. 5). Treatment also did not have sex-dependent effect on gene expression (treatment-sex interactions, p > 0.26). Females had higher expression of *NCOA1* (F=7.68, p=0.012) and *THRA* (F=5.17, p=0.034) than males.

**Figure 5.**
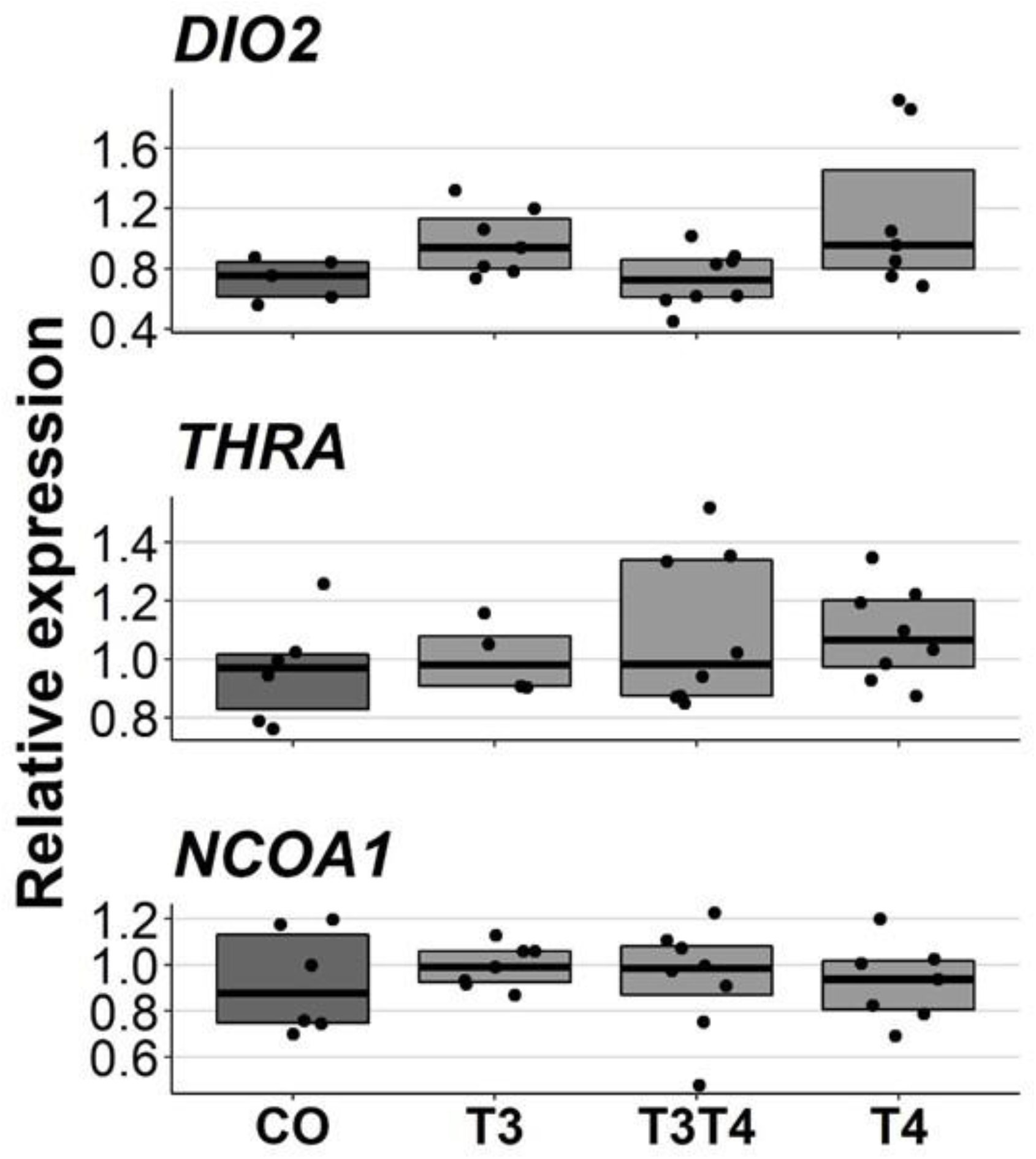
Relative expressions of DIO2, THRA and NCOA1 in the pallium of 10-month-old birds between different prenatal thyroid hormone treatments. CO = control, 50 µl 0.9% saline; T3: 4.7 ng/egg in 50 µl 0.9% saline, equivalent to 1.24 pg/mg yolk; T4: 8.9 ng/egg in 50 µl 0.9% saline, equivalent to 1.79 pg/mg yolk. The lines represent the medians of the treatments, and the bars represent the interquartile range of the original observations. Sample sizes (CO/T3/T3T4/T4): DIO2 5/7/8/7, THRA 6/4/8/8 and NCOA1 6/7/8/7.

## Discussion

In this study, we tested for long-term changes in behaviour, plasma TH levels and pallial gene expression following increased exposure to maternal THs. In terms of behaviour, we observed no significant changes in boldness, social motivation or fearfulness to predators in chicks (3-12 d) or adults (4.5 months) when the egg TH level was increased. To our knowledge, there are no similar previous studies, and it is therefore difficult to assess whether there may have been changes in other behavioural traits. The detection of changes was also partly limited by the small sample size, which meant that we may not have had enough statistical power to detect subtle changes in behaviour. Nevertheless, we found some age-dependent and sex-dependent effects of TH treatment on behaviour. The age-dependent effect in the emergence test suggests that elevated maternal TH leads to a decrease in boldness and social motivation with age (from chick to adult), whereas such a decrease is not observed in controls. However, the lack of differences between control and treatment within a given age suggests that such an effect (if not associated with type I statistical error) is rather mild. The sex-dependent effect in the tonic immobility test also seems relatively unconvincing because no sex differences were observed within treatments, nor did T3T4 females or T4 males differ from the same-sex controls. Given the important role of THs in brain development, we expected that elevated maternal THs would have led to behavioural changes. Yet, our data suggest that an experimental increase in maternal THs within the natural range has no clear and large effect on the behavioural traits measured. However, in contrast to maternal THs, post-hatch TH treatment twenty hours after hatching has been shown to affect imprinting and learning in four-day-old chicks (Yamaguchi et al., 2012). The discrepancy between our results and Yamaguchi et al. (2012) could suggest that the sensitive period during which THs can influence behaviour might occur at a later timing of development, but this requires further investigation.

We found that elevated maternal THs did not significantly affect plasma TH levels in adulthood (4.5 months of age), although transient effects prior to measurement, or downregulation of THs,cannot be ruled out. This was the first study to test for long-term changes, whereas previously changes in response to elevated prenatal THs have been observed at younger ages: Experimental elevation of egg THs was found to increase male and decrease female plasma TH levels in rock pigeons at 14 days of age (Hsu et al., 2017), but such effects were not observed in two other wild passerine species (Hsu et al., 2023b, 2020). In contrast to these studies in altricial species, not only was this study conducted on a precocial species, but our measurements were also made at a different life stage (adulthood). Overall, this suggests that an increase in maternal THs may only lead to transient changes in offspring plasma TH levels and that the effects may not be universal across species.

We also found no significant effect of elevated maternal THs on the expression of *THRA, DIO2* or *NCOA1* in the pallium of adult (10 months old) individuals. Since we only collected samples from 10-month-old birds, we cannot rule out possible changes at younger ages. Changes in gene expression have been observed in other TH-related genes, but, at much shorter time spans: In chickens, injection of THs into the yolk decreased the expression of the TH-transporter *OATP1C1* and increased the expression of the TH-transporter *MCT8* in 4-day-old embryos (Van Herck et al., 2012). The expressions of our target genes, *THRA, DIO2* and *NCOA1,* have also been shown to be at least transiently responsive to THs in other species: increase in *THRA* and decrease in *DIO2* mRNA levels in zebrafish embryos (Walpita et al., 2007), and increase in *NCOA1* mRNA levels in the anterior pituitary of adult rats (Misiti et al., 1998). It is therefore possible that the gene regulation by maternal THs is restricted to early development when the embryo is not yet able to produce THs.

We have previously shown in these same experimental birds, that an increase in prenatal THs influences hatching success (Sarraude et al., 2020). Therefore, the dose used in this study can have some effects during embryo development, which suggests that the lack of large changes in long-term behaviour, plasma TH levels and brain gene expression may be linked to an attenuation of prenatal TH effects by embryonic regulation. It is indeed possible that embryos could buffer the effect of maternal THs by altering the expression of deiodinases which are expressed already at very early stages (Ruuskanen et al., 2022; Too et al., 2017). Metabolisation of THs to inactive forms could prevent the elevated egg THs from exerting any effects. However, given the effects previously observed in these same birds on hatching success (Sarraude et al., 2020), the lack of changes in this study may also be explained by the stage of ontogeny, prenatal THs not leading to long-lasting effects. Nevertheless, the lack of prenatal measurements means that this cannot be fully confirmed, and it remains possible that elevated maternal THs have no effect on the measured traits in Japanese quails.

## Conclusions

In conclusion, we found no evidence that elevated maternal THs cause substantial long-term changes in behaviour, plasma TH levels or pallial gene expression. This is consistent with our previous findings in the same experimental birds, where elevation of prenatal THs (T3, T4, or in combination) did not lead to long-term effects on oxidative stress, moult or reproduction (Sarraude et al., 2020). While behavioural changes have not been previously studied, plasma TH concentrations and gene expression have been shown to respond to elevated maternal THs in the short-term (Hsu et al., 2017; Van Herck et al., 2012). However, based on this study, these changes would appear to be limited to early development or shortly after hatching with the offspring’s own TH production later replacing the effects of elevated maternal THs. Maternal THs could theoretically influence the offspring’s own TH production at a later age by altering the function of the HPT-axis, but no evidence has yet been found to support this hypothesis. The long-term effects of variation in maternal THs remain unclear, but this study has provided new information on some previously untested traits. Given the tight association between THs and metabolism, it would be interesting to study behaviors related to thermoregulation and metabolism in the future, Understanding long-term effects in addition to short-term effects will help to better understand the evolutionary and ecological importance of maternal THs.

## Acknowledgements

We thank personnel at the animal care facilities, Marjo Myllyrinne, Simo Laine for their help.

## Data availability statement

Data is available on Figshare (DOI: https://doi.org/10.6084/m9.figshare.22266211.v1).

---

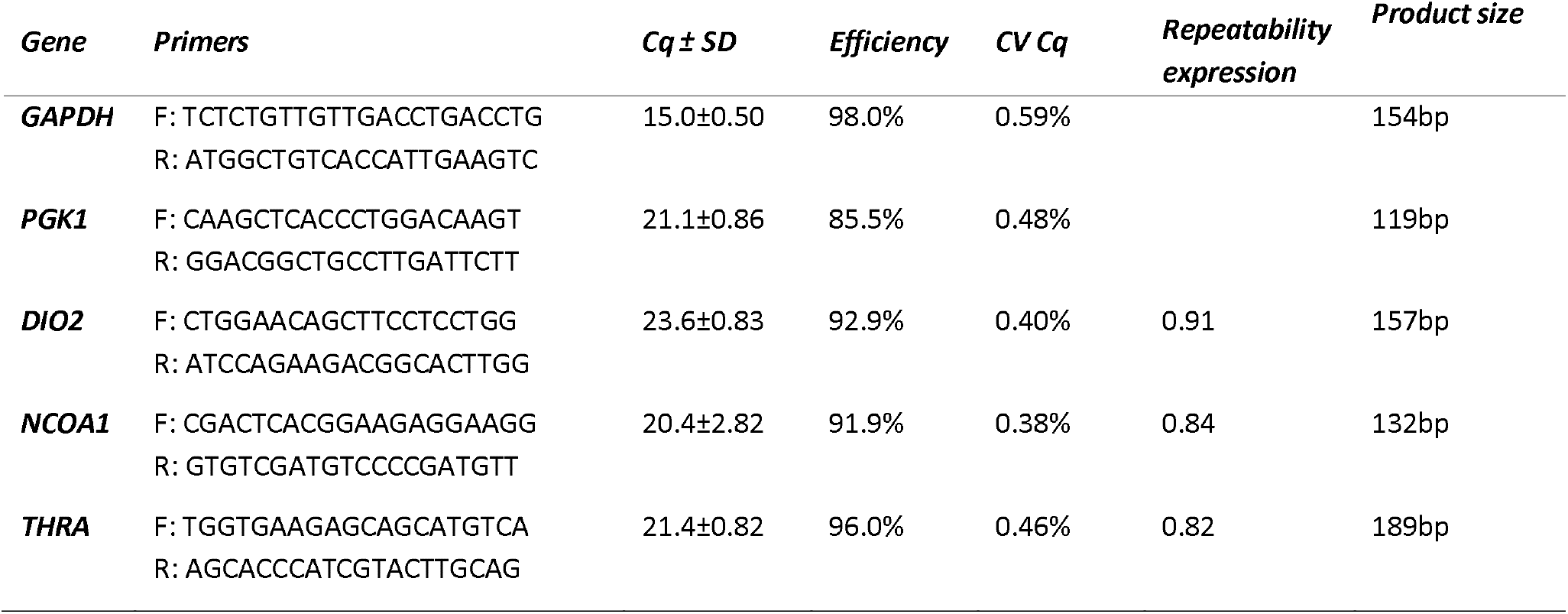

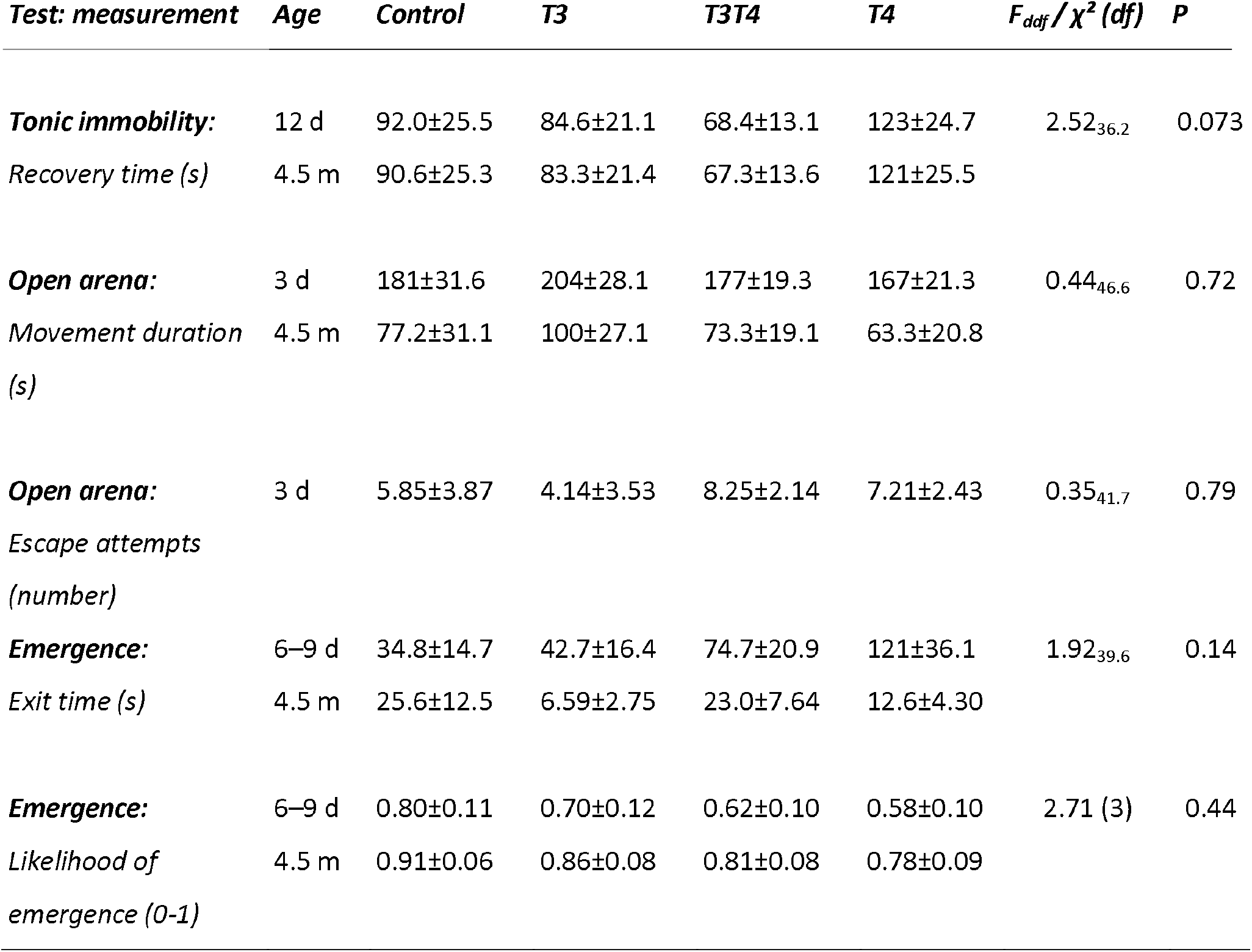

| *Test: measurement* | *Age* | *Control* | *T3* | *T3T4* | *T4* | *F<sub>ddf</sub>/χ<sup>2</sup> (df)* | *P* |
| --- | --- | --- | --- | --- | --- | --- | --- |
| ***Tonic immobility:*** | 12 d | 92.0±25.5 | 84.6±21.1 | 68.4±13.1 | 123±24.7 | 2.52<sub>36,2</sub> | 0.073 |
| *Recovery time (s)* | 4.5 m | 90.6±25.3 | 83.3±21.4 | 67.3±13.6 | 121±25.5 |  |  |
| ***Open arena:*** | 3 d | 181±31.6 | 204±28.1 | 177±19.3 | 167±21.3 | 0.44<sub>46,6</sub> | 0.72 |
| *Movement duration (s)* | 4.5 m | 77.2±31.1 | 100±27.1 | 73.3±19.1 | 63.3±20.8 |  |  |
| ***Open arena:*** | 3 d | 5.85±3.87 | 4.14±3.53 | 8.25±2.14 | 7.21±2.43 | 0.35<sub>41,7</sub> | 0.79 |
| *Escape attempts (number)* |  |  |  |  |  |  |  |
| ***Emergence:*** | 6–9 d | 34.8±14.7 | 42.7±16.4 | 74.7±20.9 | 121±36.1 | 1.92<sub>39,6</sub> | 0.14 |
| *Exit time (s)* | 4.5 m | 25.6±12.5 | 6.59±2.75 | 23.0±7.64 | 12.6±4.30 |  |  |
| ***Emergence:*** | 6–9 d | 0.80±0.11 | 0.70±0.12 | 0.62±0.10 | 0.58±0.10 | 2.71 (3) | 0.44 |
| *Likelihood of emergence (0-1)* | 4.5 m | 0.91±0.06 | 0.86±0.08 | 0.81±0.08 | 0.78±0.09 |  |  |

