## Supplemental figure 1 for "Can increased prenatal exposure to thyroid hormones alter physiology and behaviour in the long-term? Insights from an experimental study in Japanese quails"

Tonic immobility

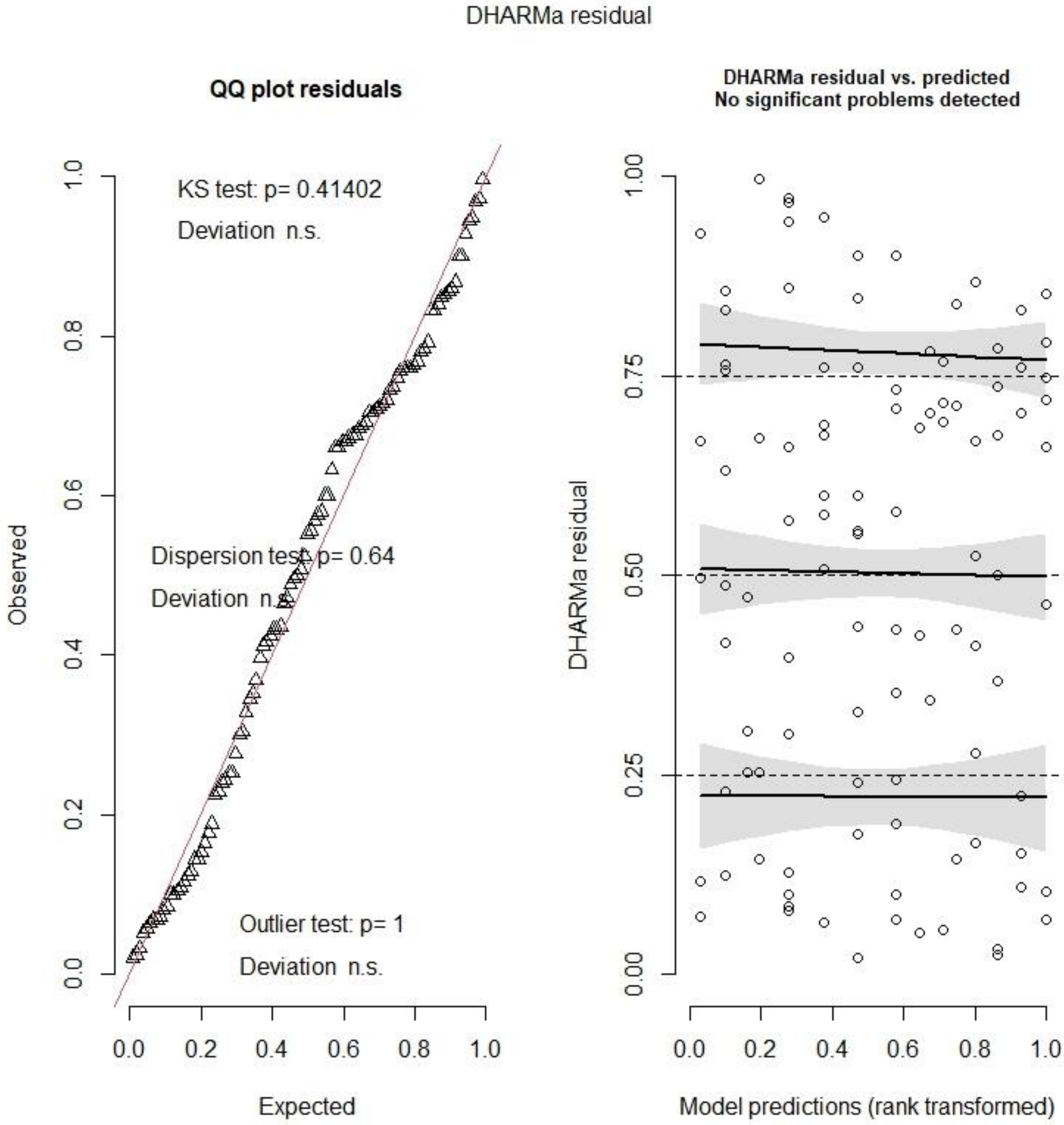

Open arena, chick escapes

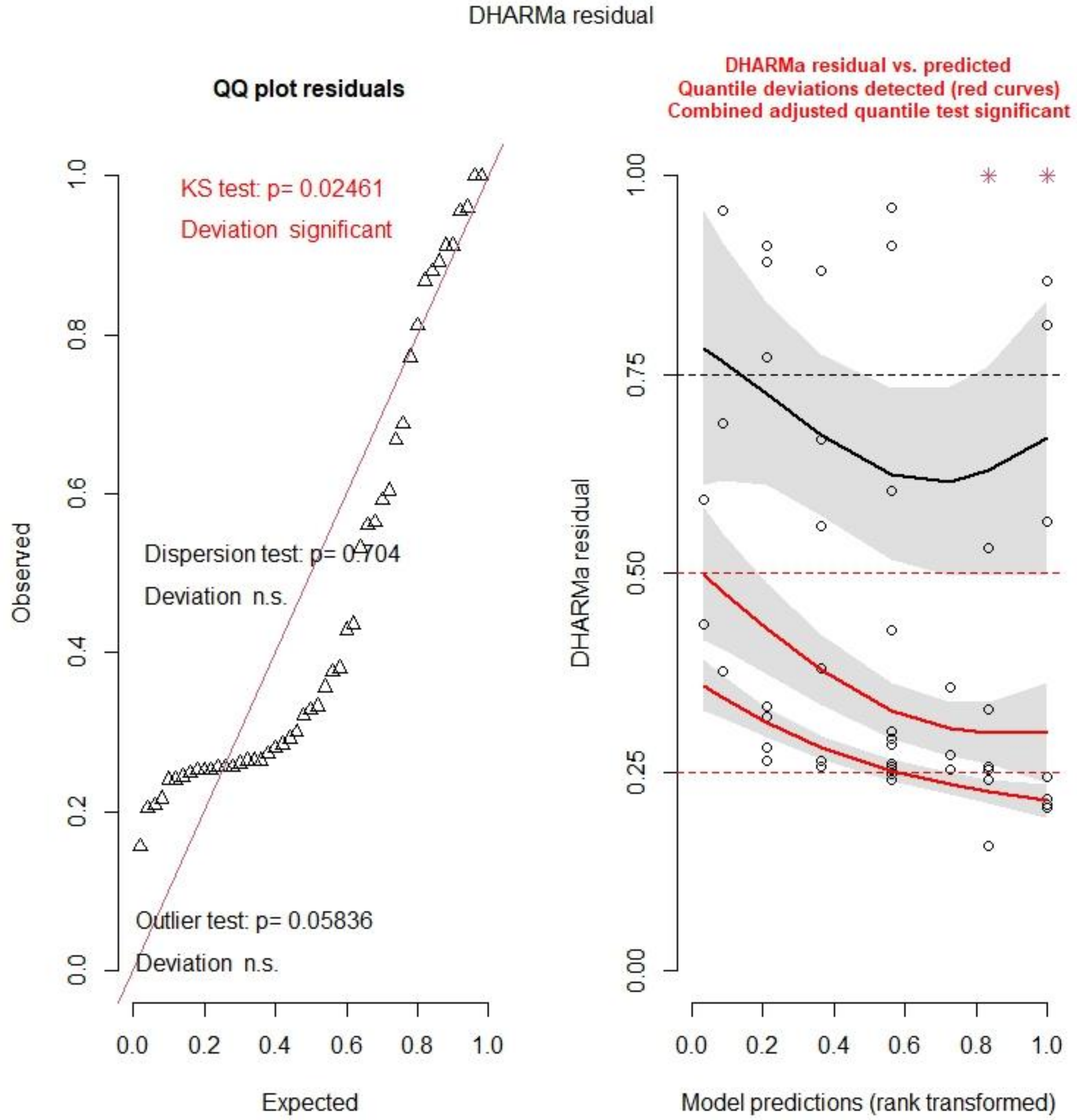

Open arena, movement duration

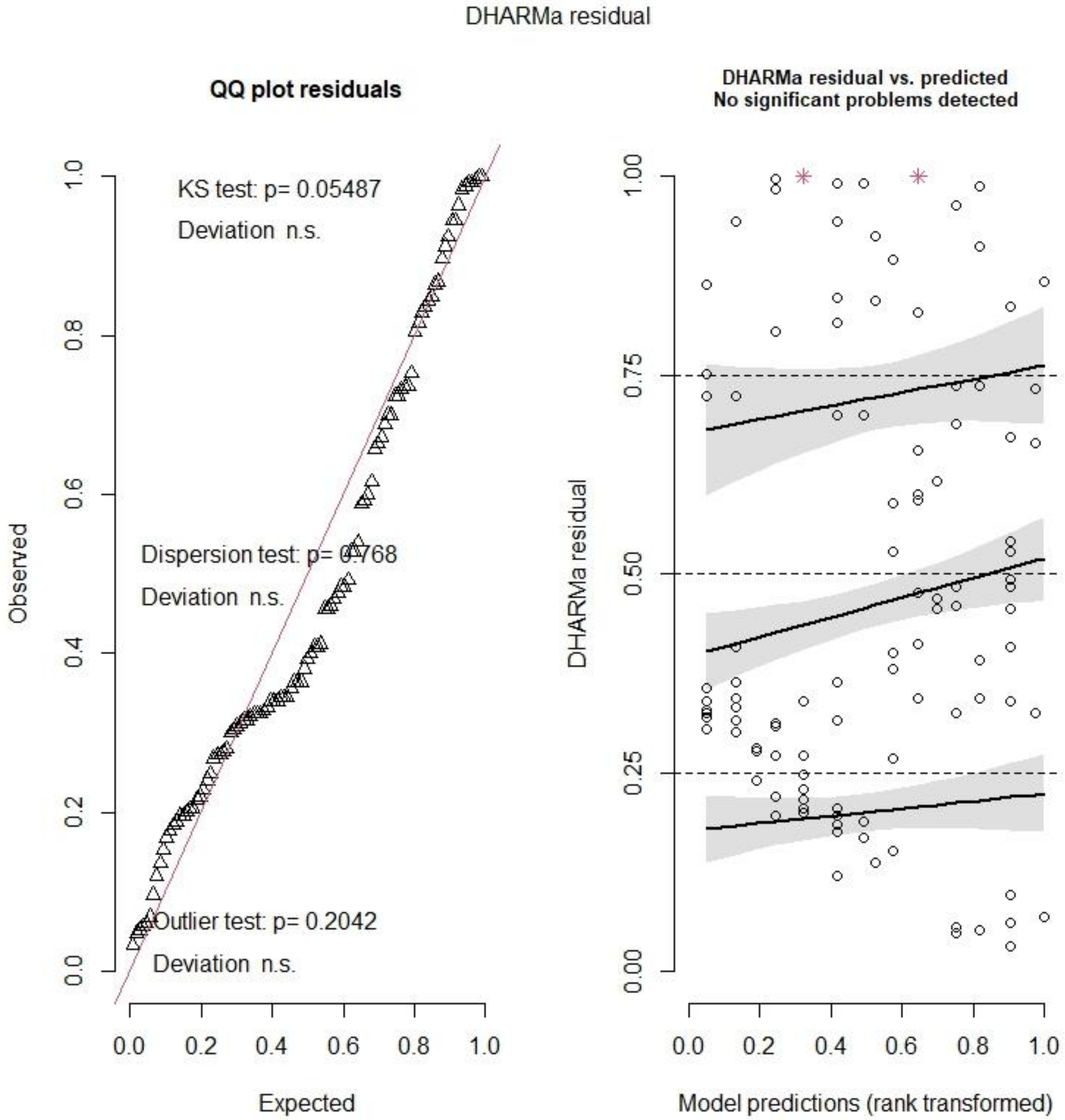

Emergence test, time until exit

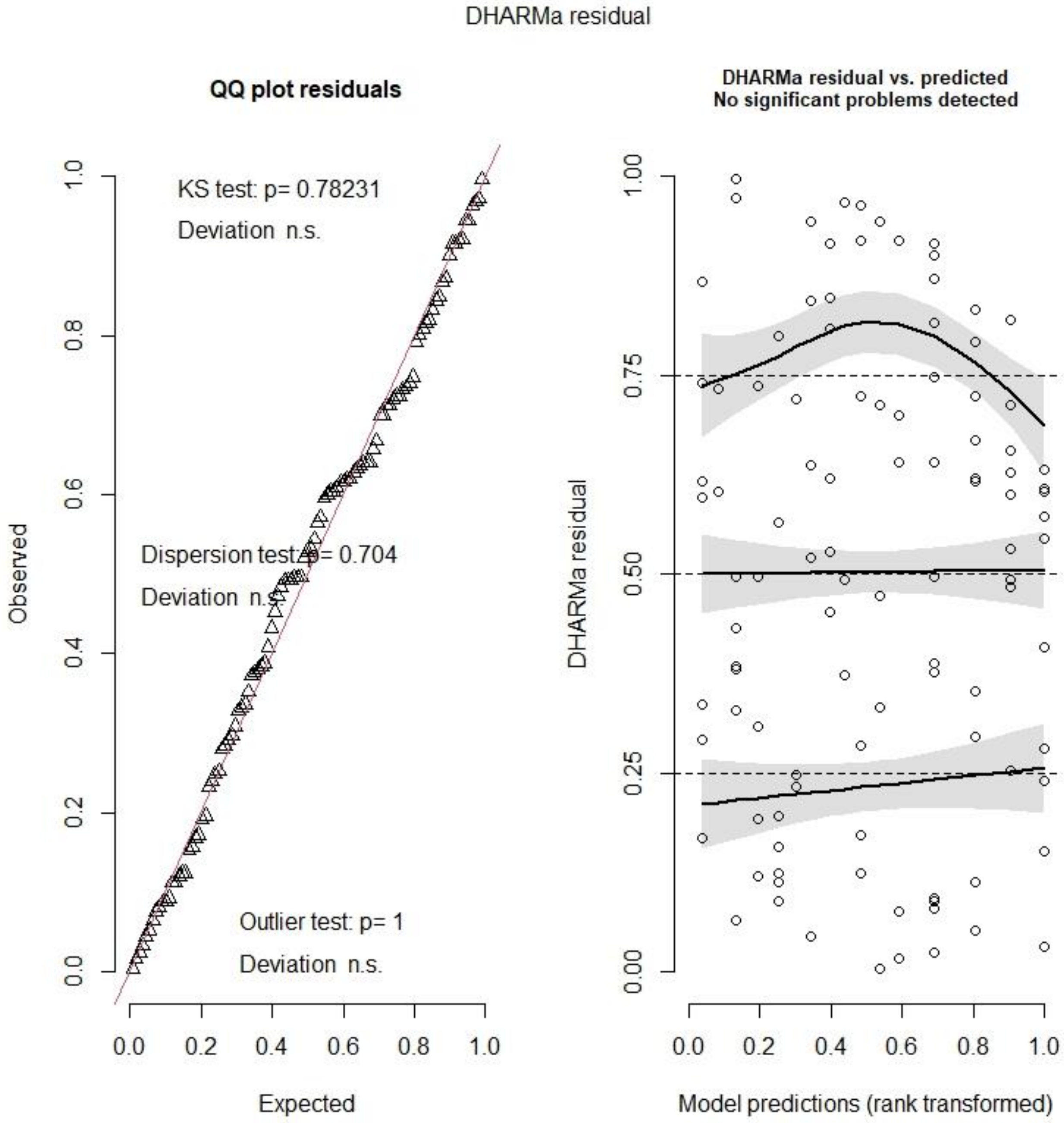

Emergence test, likelihood of emergence

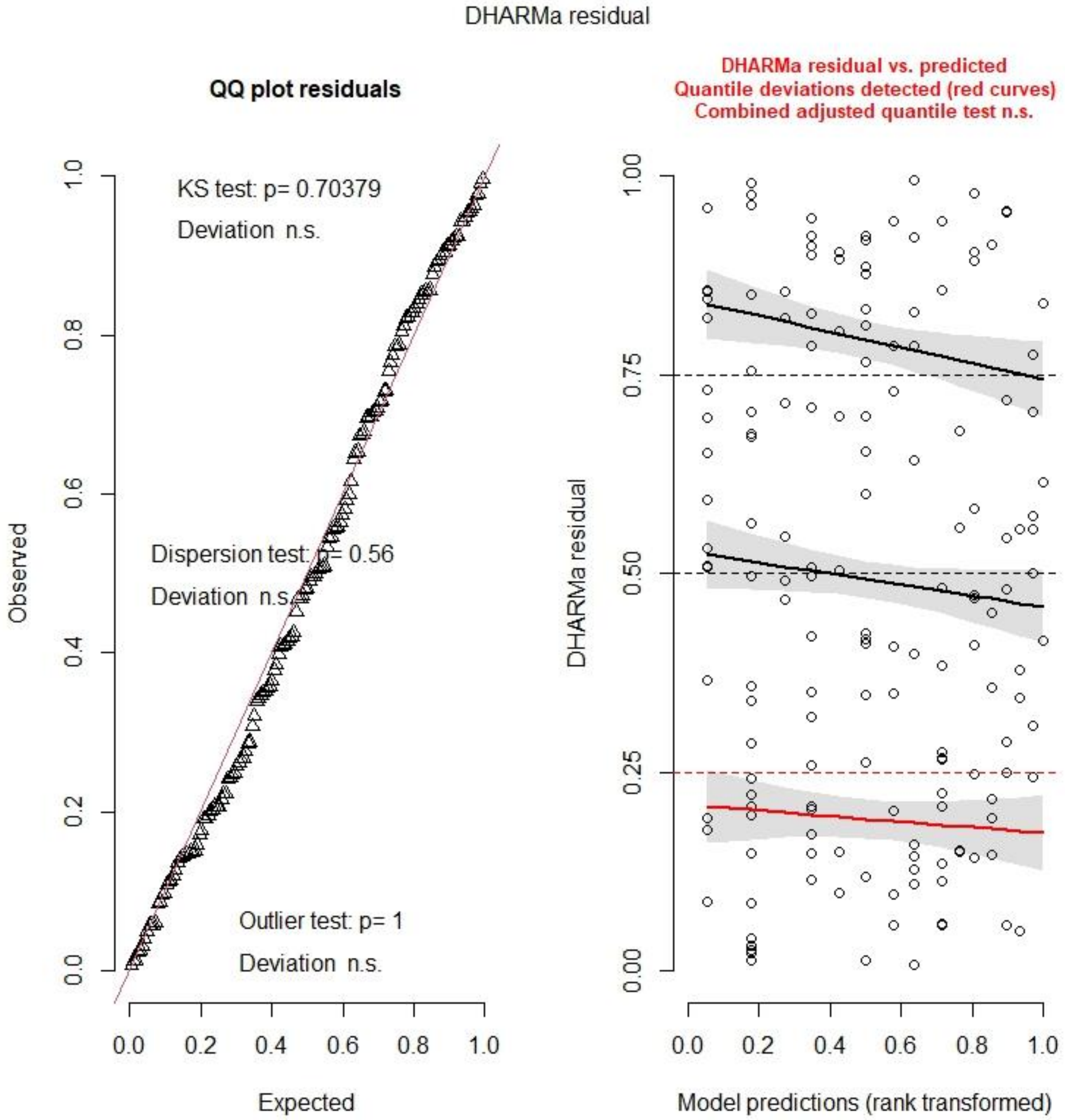

T3 concentration

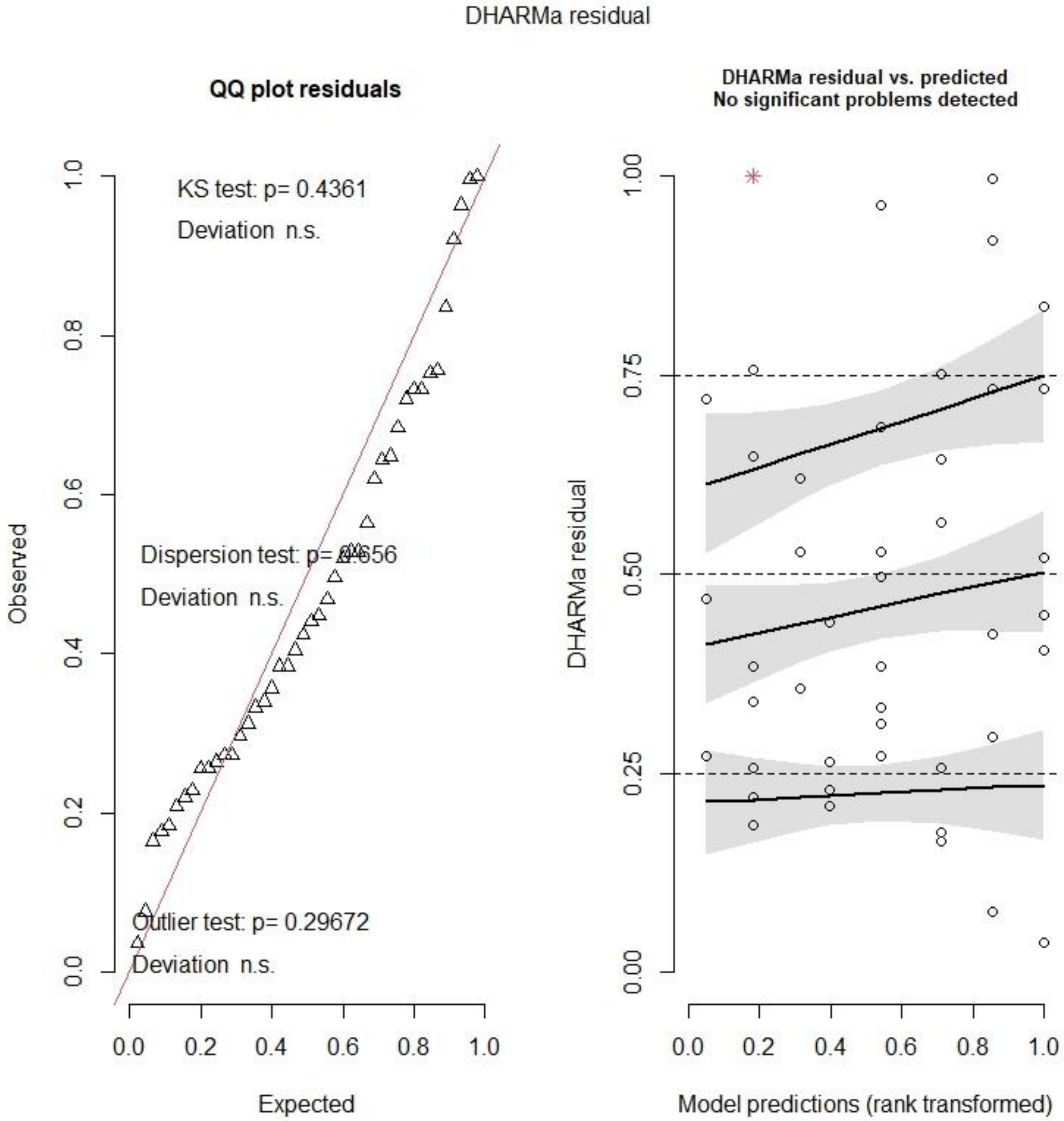

T4 concentration

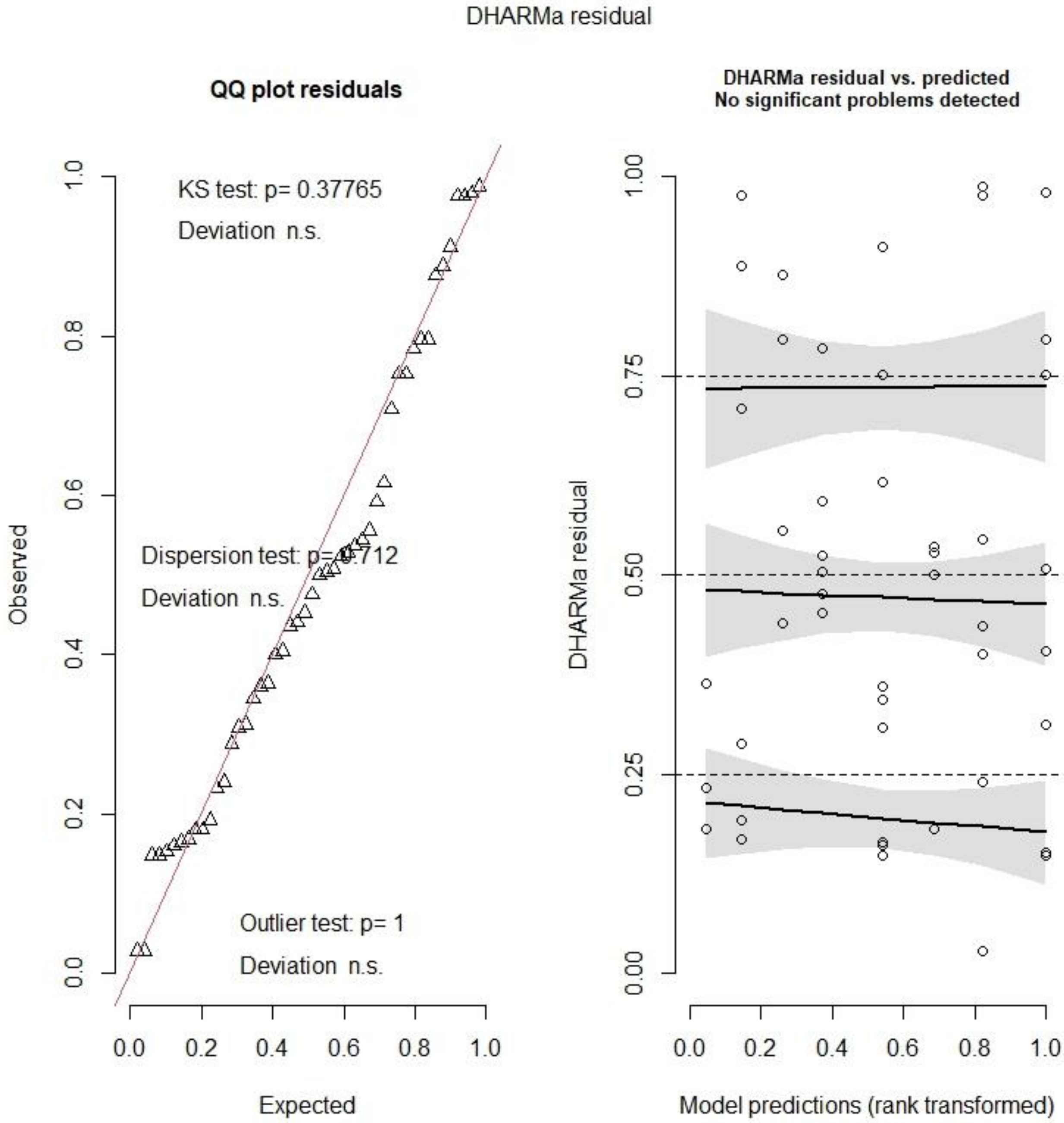

THRA expression

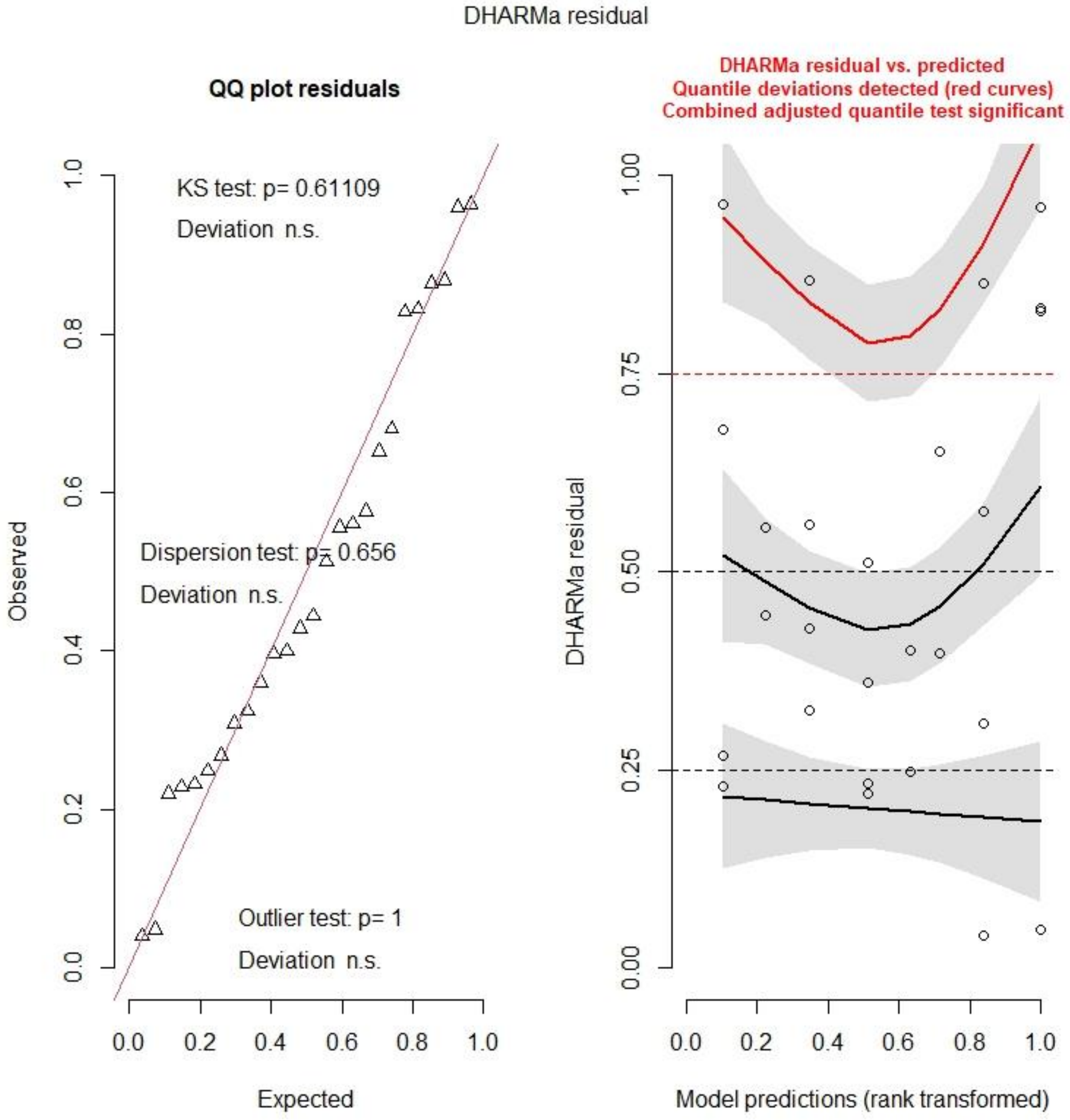

NCOA1 expression

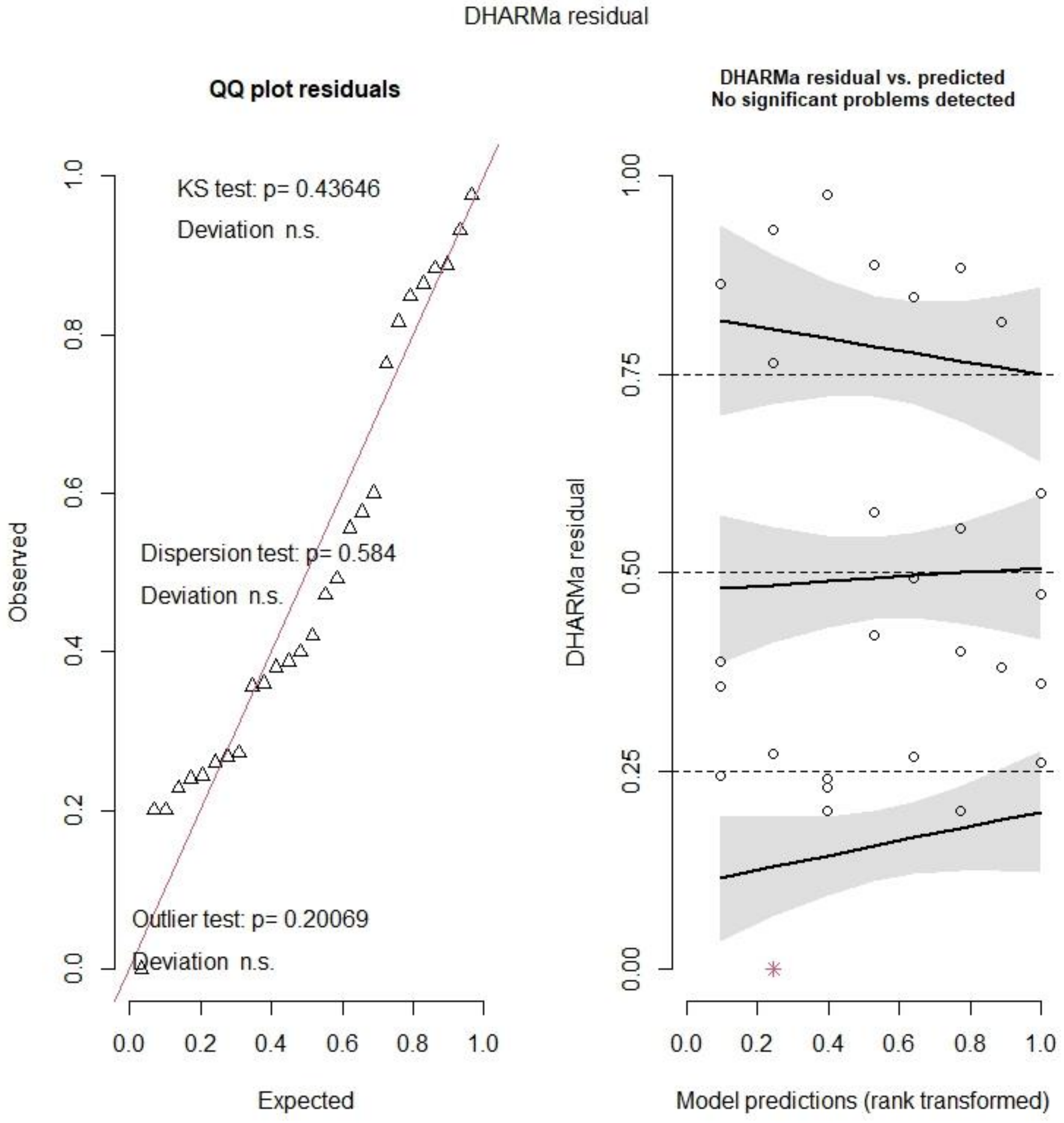

DIO2 expression

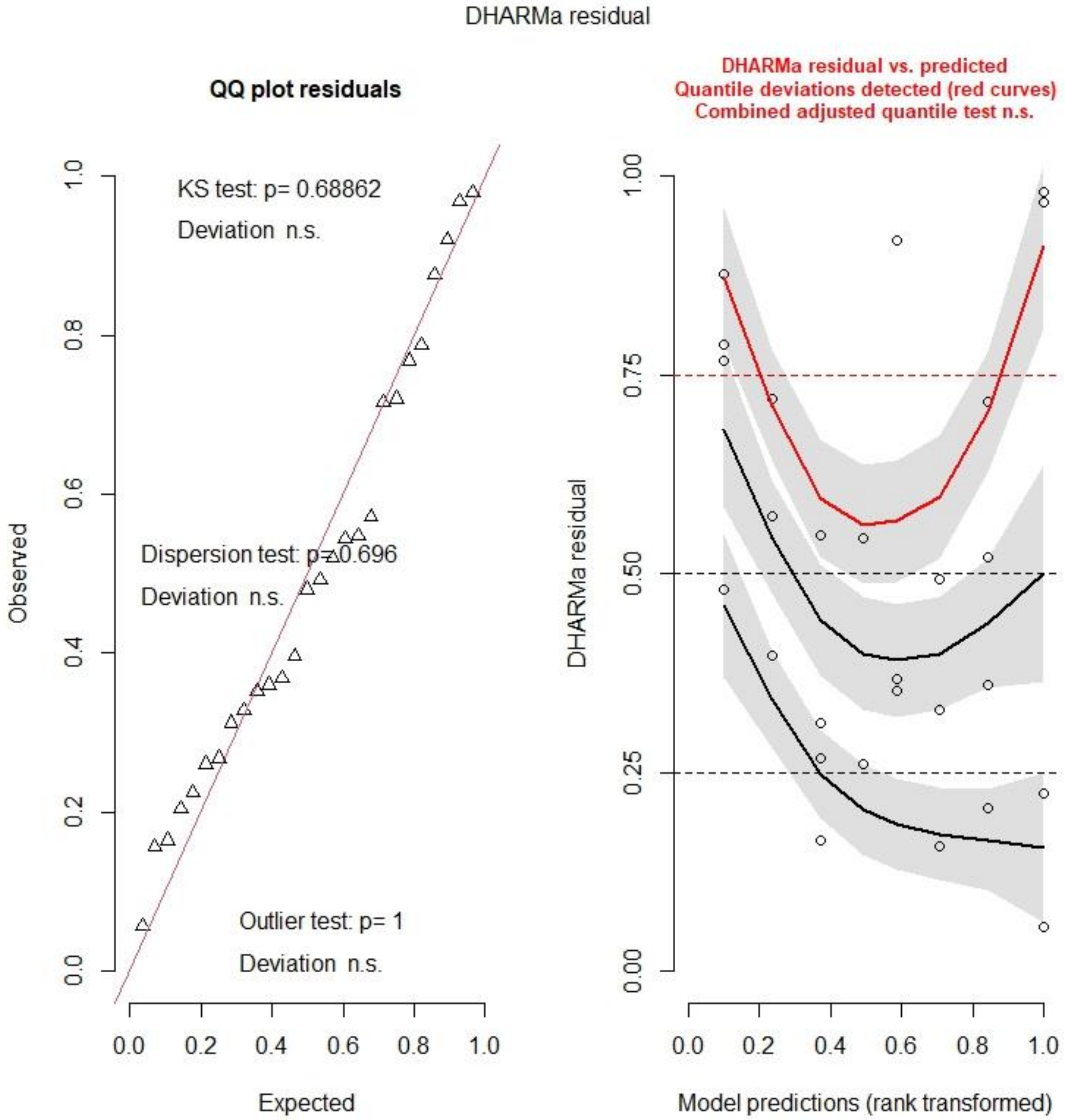
