## Supplemental table 1 for "Can increased prenatal exposure to thyroid hormones alter physiology and behaviour in the long-term? Insights from an experimental study in Japanese quails"

| **Test, measurement** | **Age** | **Mean (Standard error)** | | | |
| --- | --- | --- | --- | --- | --- |
|  |  | *Control* | *T3* | *T3T4* | *T4* |
| Tonic Immobility,  *Recovery time (s)* | Chicks (12 days) | 126.4 (26.7) | 141.3 (45.2) | 95.6 (16.1) | 159.8 (30.9) |
|  | Adults (4.5 months) | 103.4 (25.2) | 110.3 (32.8) | 131.7 (35.0) | 168.9 (35.1) |
| Open Arena,  *Escape attempts* | Chicks (3 days) | 5.0 (2.9) | 4.9 (1.8) | 7.9 (2.2) | 6.9 (2.4) |
| *Movement duration (s)* | Chicks (3 days) | 175.7 (28.0) | 215.0 (40.3) | 180.5 (19.8) | 156.3 (22.2) |
|  | Adults (4.5 months) | 65.4 (46.1) | 83.8 (39.5) | 76.7 (20.8) | 71.5 (23.8) |
| Emergence,  *Time until exit (s)* | Chicks (6–9 days) | 83.6 (28.2) | 86.6 (23.7) | 116.5 (19.9) | 136.7 (16.0) |
|  | Adults (4.5 months) | 58.5 (36.9) | 9.1 (2.1) | 41.9 (12.9) | 32.6 (17.0) |
| Plasma T3 concentration (pmol/ml) | Adults (4.5 months) | 4.44 (0.46) | 5.25 (0.91) | 4.96 (0.76) | 5.14 (0.54) |
| Plasma T4 concentration (pmol/ml) |  | 6.02 (1.52) | 7.57 (0.45) | 7.12 (0.97) | 9.95 (1.31) |
| Brain DIO2 expression | Adults (10 months) | 0.73 (0.06) | 0.98 (0.08) | 0.73 (0.07) | 1.15 (0.20) |
| Brain NCOA1 expression |  | 0.93 (0.09) | 1.00 (0.04) | 0.94 (0.08) | 0.93 (0.06) |
| Brain THRA expression |  | 0.96 (0.07) | 1.01 (0.06) | 1.10 (0.09) | 1.09 (0.06) |
