## Supplemental table 2 for "Can increased prenatal exposure to thyroid hormones alter physiology and behaviour in the long-term? Insights from an experimental study in Japanese quails"

|  | **Tonic Immobility,**  **Recovery time** | | **Open Arena,**  **Escape attempts** | | **Open Arena,**  **Movement duration** | | | **Emergence,**  **Time until exit** | | **Emergence,**  **Likelihood of emergence** | | **Plasma**  **T3 concentration** | | **Plasma**  **T4 concentration** | | **Brain**  **DIO2 expression** | | **Brain**  **NCOA1 expression** | | **Brain**  **THRA**  **expression** | |
| --- | --- | --- | --- | --- | --- | --- | --- | --- | --- | --- | --- | --- | --- | --- | --- | --- | --- | --- | --- | --- | --- |
| **N Chicks** (F/M) Control, T3, T3T4, T4 | 3/4, 7/4, 9/11, 10/8 | | 3/3, 6/3, 8/12, 8/8 | | | 3/3, 6/3, 8/12, 8/8 | | 5/5, 6/7, 14/10, 12/10 | | 6/8, 13/8, 17/24, 20/16 | |  | | | |  | | | | | |
| **N Adults** | 3/4, 6/3, 5/12, 8/6 | | - | | 3/4, 8/3, 9/12, 9/8 | | | 3/3, 7/2, 7/6, 8/5 | | 3/4, 7/4, 8/11, 9/7 | | 3/3, 6/4, 6/8, 6/8 | | 3/4, 6/4, 6/9, 8/8 | | 3/2, 3/4, 4/4, 3/4 | | 2/4, 4/3, 4/4, 3/4 | | 2/4, 2/2, 4/4, 4/4 | |
| **N Mothers** | 21 | | 19 | | | | | 21 | | | | 19 | | 20 | | 17 | | 16 | | 16 | |
| **Predictor** | Estimate (SE) | P | Estimate (SE) | P | Estimate (SE) | | P | Estimate (SE) | P | Estimate (SE) | P | Estimate (SE) | P | Estimate (SE) | P | Estimate (SE) | P | Estimate (SE) | P | Estimate (SE) | P |
| T3 | 0.36  (0.45) | 0.42 | -1.70  (4.97) | 0.73 | 23.25 (38.16) | | 0.55 | -1.36 (0.62) | 0.03 | -0.57  (0.77) | 0.46 | 0.75  (1.27) | 0.56 | 1.86  (1.98) | 0.35 | 0.22 (0.18) | 0.22 | 0.02  (0.09) | 0.83 | 0.01  (0.12) | 0.90 |
| T3T4 | -0.47  (0.43) | 0.28 | 2.41  (4.24) | 0.57 | -3.88 (33.34) | | 0.91 | -0.11 (0.57) | 0.85 | -0.92  (0.69) | 0.18 | 0.58  (1.20) | 0.64 | 1.04  (1.83) | 0.57 | -0.02 (0.16) | 0.88 | -0.02  (0.09) | 0.81 | 0.11  (0.10) | 0.30 |
| T4 | -0.02  (0.43) | 0.96 | 1.37  (4.40) | 0.76 | -13.81 (34.41) | | 0.69 | -0.71 (0.58) | 0.22 | -1.11 (0.74) | 0.13 | 0.75  (1.20) | 0.54 | 4.05  (1.82) | 0.03 | 0.37 (0.18) | 0.05 | -0.02  (0.09) | 0.81 | 0.09  (0.10) | 0.35 |
| Sex (male) | 0.09  (0.50) | 0.86 | -3.19 (2.71) | 0.24 | 36.52 (21.01) | | 0.09 | 0.43 (0.28) | 0.13 | -0.83  (0.45) | 0.07 | -0.69 (0.75) | 0.36 | 1.80  (1.17) | 0.13 | 0.17 (0.12) | 0.18 | -0.19  (0.06) | 0.01 | -0.18  (0.07) | 0.02 |
| Age (chick) | 0.02  (0.17) | 0.92 | - | | 103.98 (18.42) | | <0.001 | 0.31 (0.52) | 0.56 | -0.96 (0.43) | 0.02 | - | | | | - | | | | | |
| % Variance explained by parental ID | 21.3 % | | 3.9 % | | 0 % | | | 15 % | | - | | 0 % | | 0 % | | 39.4 % | | 0 % | | 0 % | |
